# Sequence effects on size, shape, and structural heterogeneity in Intrinsically Disordered Proteins

**DOI:** 10.1101/427476

**Authors:** Upayan Baul, Debayan Chakraborty, Mauro L. Mugnai, John E. Straub, D. Thirumalai

**Affiliations:** Department of Chemistry, The University of Texas at Austin, Austin, Texas 78712; Department of Chemistry, Boston University, Boston, Massachusetts 02215; Institute of Physics, Albert-Ludwigs-University of Freiburg, Hermann-Herder-Strasse 3, 79104 Freiburg, Germany

## Abstract

Intrinsically disordered proteins (IDPs) lack well-defined three-dimensional structures, thus challenging the archetypal notion of structure-function relationships. Determining the ensemble of conformations that IDPs explore under physiological conditions is the first step towards understanding their diverse cellular functions. Here, we quantitatively characterize the structural features of IDPs as a function of sequence and length using coarse-grained simulations. For diverse IDP sequences, with the number of residues (*N*_*T*_) ranging from 24 to 441, our simulations not only reproduce the radii of gyration (*R*_*g*_) obtained from experiments, but also predict the full scattering intensity profiles in very good agreement with Small Angle X-ray Scattering experiments. The *R*_*g*_ values are well-described by the standard Flory scaling law, 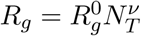, with *v* ≈ 0.588, making it tempting to assert that IDPs behave as polymers in a good solvent. However, clustering analysis reveals that the menagerie of structures explored by IDPs is diverse, with the extent of heterogeneity being highly sequence-dependent, even though ensemble-averaged properties, such as the dependence of *R*_*g*_ on chain length, may suggest synthetic polymer-like behavior in a good solvent. For example, we show that for the highly charged Prothymosin-*α*, a substantial fraction of conformations is highly compact. Even if the sequence compositions are similar, as is the case for *α*-Synuclein and a truncated construct from the Tau protein, there are substantial differences in the conformational heterogeneity. Taken together, these observations imply that metrics based on net charge or related quantities alone, cannot be used to anticipate the phases of IDPs, either in isolation or in complex with partner IDPs or RNA. Our work sets the stage for probing the interactions of IDPs with each other, with folded protein domains, or with partner RNAs, which are critical for describing the structures of stress granules and biomolecular condensates with important cellular functions.

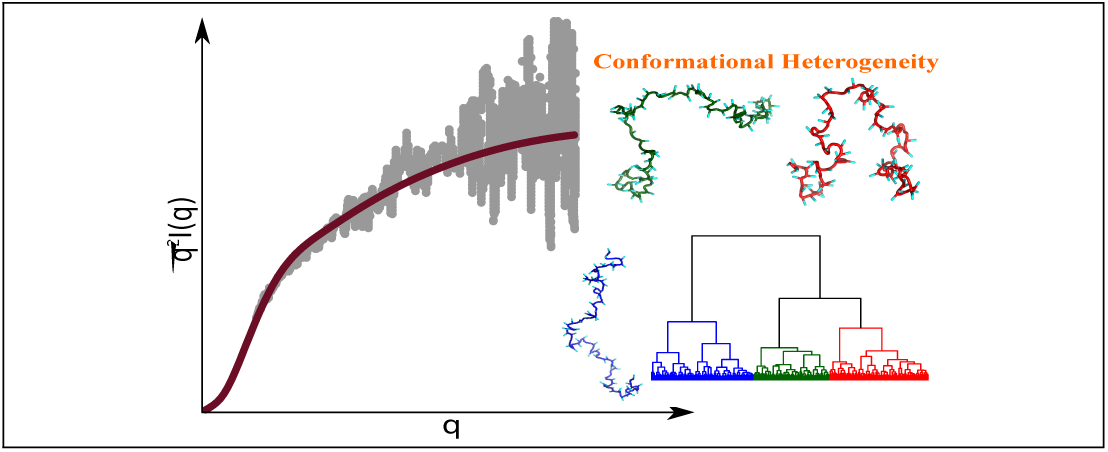
Graphical TOC Entry.

## Introduction

The discovery that a large fraction of eukaryotic protein sequences are not ordered in isolation^1–7^ has produced a paradigm shift in the commonly held view of structure-function relationship. Such sequences, referred to as Intrinsically Disordered Proteins (IDPs), play key functional roles in diverse cellular processes, such as signal transduction, ^2,8^ and vesicular transport,^9^ and are also implicated in neurodegenerative disorders and other diseases.^3,10^ Experimental realizations of folding coupled to binding partners, and the more recent discovery of intracellular liquid-liquid phase separation^11^ have further resulted in a concerted effort in describing the intramolecular and intermolecular interactions involving IDPs.^12–17^ Recent estimates indicate that between (30-40)% of eukaryotic proteomes are either intrinsically disordered, or contain intrinsically disordered regions in otherwise folded proteins.^17,18^ Despite the importance of IDPs, characterization of the sequence-dependent conformational properties of IDPs in isolation, as well as their phase behavior remains largely qualitative.^14,16,19^ Biologically relevant forms of IDPs do not fold into unique tertiary structures, but explore a large number of distinct conformational states. In this sense, they are like synthetic polymers, whose conformations can only be characterized in statistical terms, such as the distribution of the radius of gyration, *R*_*g*_, from which its dependence on the number of residues, *N*_*T*_, can be calculated.^17,20^ Two decades of experiments indicate that isolated IDPs typically behave as polymers in a good solvent, based on the scaling of *R*_*g*_ with *N*_*T*_.^21–23^ However, the conformational ensembles of IDPs, which are instrumental in modulating their *in vivo* functionalities, strongly depend on the precise sequence, in addition to other external conditions, such as pH, temperature, and salt concentration.^24,25^ Therefore, scaling laws alone cannot provide a faithful description of the physico-chemical properties of IDPs. The complex sequence dependence poses a great challenge to purely theoretical approaches,^26,27^ and raises the need for computational models that can quantitatively account for the experimental data.

Small-angle X-ray scattering (SAXS) and single-molecule Förster resonance energy transfer (smFRET) have found the greatest applicability in the study of IDP conformational properties.^21,28^ In addition, Nuclear Magnetic Resonance (NMR) and fluorescence correlation spectroscopy (FCS)^24,28–31^ have been used to characterize the structural ensembles, and conformational dynamics of IDPs. SAXS experiments, which measure structure factors, could be used to calculate the global conformational properties of IDPs, such as *R*_*g*_ and shape, while smFRET experiments are used to probe distances between specific labeled residues along the polypeptide chain. With SAXS, the measured form factor, *I*(*q*), as function of the wave vector *q*, at small values of *q* is often used to estimate *R*_*g*_ using the Guinier approximation.^32^ However, the sequence-specific conformational ensembles are not directly available from SAXS experiments, and usually techniques based on ensemble-optimization are invoked to map scattering profiles to representative structures.^33^

A complementary approach involves force-field based computer simulations to generate IDP ensembles, which embody the key experimental observables. It is tempting to exploit an all-atom representation for IDPs and the surrounding environment in order to characterize their conformations. Indeed, great progress is being made in devising new force fields, which make systematic updates to the existing water models or protein interaction potentials.^34–38^ Systematic benchmarking has revealed that although all-atom force fields provide the much needed microscopic insight into the conformational dynamics of IDPs, the associated ensembles often tend to depend on the details of the parametrization.^39,40^ A simpler description of the protein molecule based on systematic coarse-graining, often provides a complementary route towards probing the conformational dynamics if IDPs.^15,41,42^ We and others have previously exploited coarse-graining strategies with great success in applications of protein folding and kinetics.^43–45^ In this work, we introduce a self-organized polymer (SOP) coarse-grained model for IDPs (SOP-IDP), which not only recapitulates the wealth of experimental SAXS data, but also delineates the complex interplay between sequence, structure, and aspects of conformational heterogeneity that have been largely unexplored in much of the previous studies.

We show that for IDPs of varying lengths, and sequence composition, the *R*_*g*_ values are in accord with Flory’s scaling law. The calculated *R*_*g*_ values from simulations are generally in good agreement with those extracted from SAXS data. The simulations accurately reproduce the measured SAXS profiles, thus allowing us to provide insights into the shape, and conformational fluctuations, which govern their functions. The ensembles of conformations of all the IDPs are heterogeneous, with sequence being a key determinant of the relative populations of different substates. Our findings using SOP-IDP simulations suggest that the phases of IDPs cannot be predicted solely based on sequence-compositional properties, but requires complete statistical characterization of the IDP ensembles.

## Results

### SOP-IDP model accurately reproduces experimental SAXS profiles

We used the SOP-IDP model (see Methods and SI for details) to calculate a variety of measurable structural properties for different IDP sequences. Twelve of these, including the initial training set consisting of Histatin-5, ACTR and hNHE1, are sequences with lengths ranging from 24 to 273 that are unrelated in composition and biological functions.^46–57^ The computed scattering profiles, *I*(*q*) as a function of the scattering vector *q*, bear close resemblance to those obtained from SAXS experiments, particularly in the low *q* regime, which describes the global structure of IDPs. The scattering profiles depicted in the Kratky representation (*q*^2^*I*(*q*)*/I*(0) vs *q*, Figs S1-S2) are also accurate at low *q* values. The level of agreement between our simulations and experiment is further quantified by calculating the normalized squared deviation, *δ*^2^ (Table S2). The *δ*^2^ values conclusively show that for most sequences, the overlap between the calculated and experimental SAXS profiles remains good up to *qR*_*g*_ ≈ 3, well beyond the range of validity of the Guinier approximation, normally taken to be *qR*_*g*_ ≈ 1.3. Hence, the model not only accurately describes the global shapes of IDPs, but also likely captures the structural details on smaller length scales. In some cases, the agreement with experimental profiles deteriorates at *qR*_*g*_ > 3, perhaps owing to a combined effect of the increased experimental noise, and the coarse-grained nature of the model.

### Wild-type Tau proteins and its variants

In notable SAXS experiments, Svergun and coworkers measured the form factors of the Wild-type (WT) 441 residue human Tau protein, as well as artificial sequence constructs generated through truncation and merger of segments.^58^ We simulated eleven such artificial constructs (SI contains the sequences), in addition to the full length WT sequence using the SOP-IDP model, thus covering the full range of sequence lengths (99 ≤ *N*_*T*_ ≤ 441) considered in the experiments.^58^ The scattering profiles for the Tau sequences are shown in Figure 2.

As is evident from the *δ*^2^ values (Table S2), the simulated and experimental profiles are in good agreement. It should be emphasized that we used only five (Histatin-5, ACTR, hNHE1, K32, and hTau40) of the twenty IDPs shown in Figures 1 and 2 to learn the parameters in the model (described in the SI). Of these, we used only the *R*_*g*_ values for K32 and hTau40 as a constraint to determine a single parameter, and not the SAXS profiles. Therefore, the good agreement between simulated and experimental SAXS profiles, across the sample set, which is diverse in terms of sequence composition, length and charge densities, is truly an emergent property of our model. In addition, we also show in the SI that the calculated *I*(*q*) as a function of *q* for the 24-residue RS peptide is in excellent agreement with experiments as are the results based on one of the recently introduced atomically detailed force field.^38^ Due to its predictive power over a wide range of sequence space, we anticipate that the SOP-IDP model will be efficacious in applications that require a faithful description of conformational ensembles, among other properties. We have predicted *I*(*q*) for two other IDPs in the SI, which await future validation.

**Figure 1:**
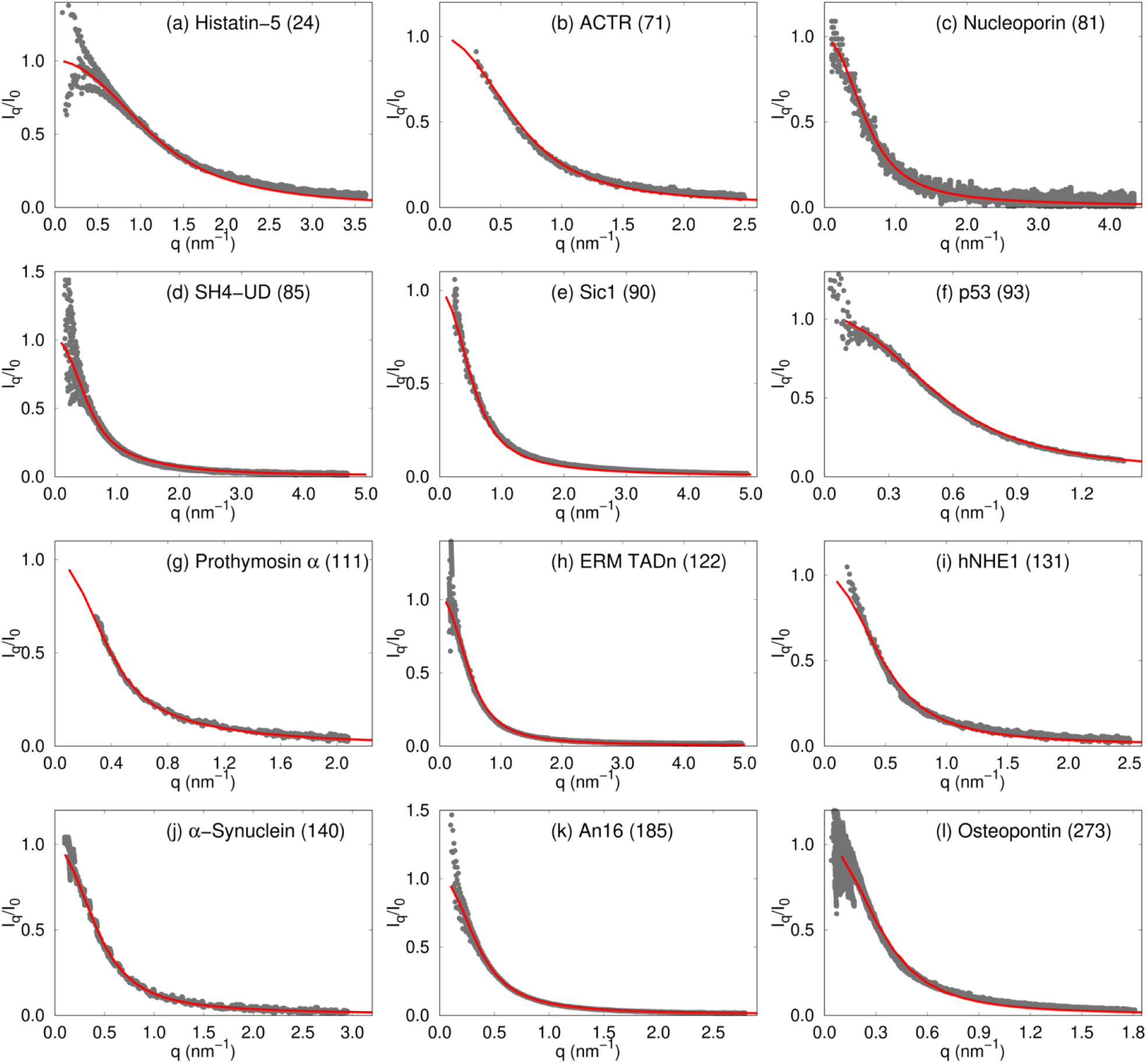
Comparison of SAXS profiles for 12 IDPs, labeled in the panels. The experimental profiles^46–56^ are shown using gray dots, and those obtained from simulations are depicted using red curves. The numbers in parentheses in each figure is *N*_*T*_, the number of residues in each IDP. Histatin-5 (a), ACTR (b) and hNHE1 (i) are part of the initial training set.

**Figure 2:**
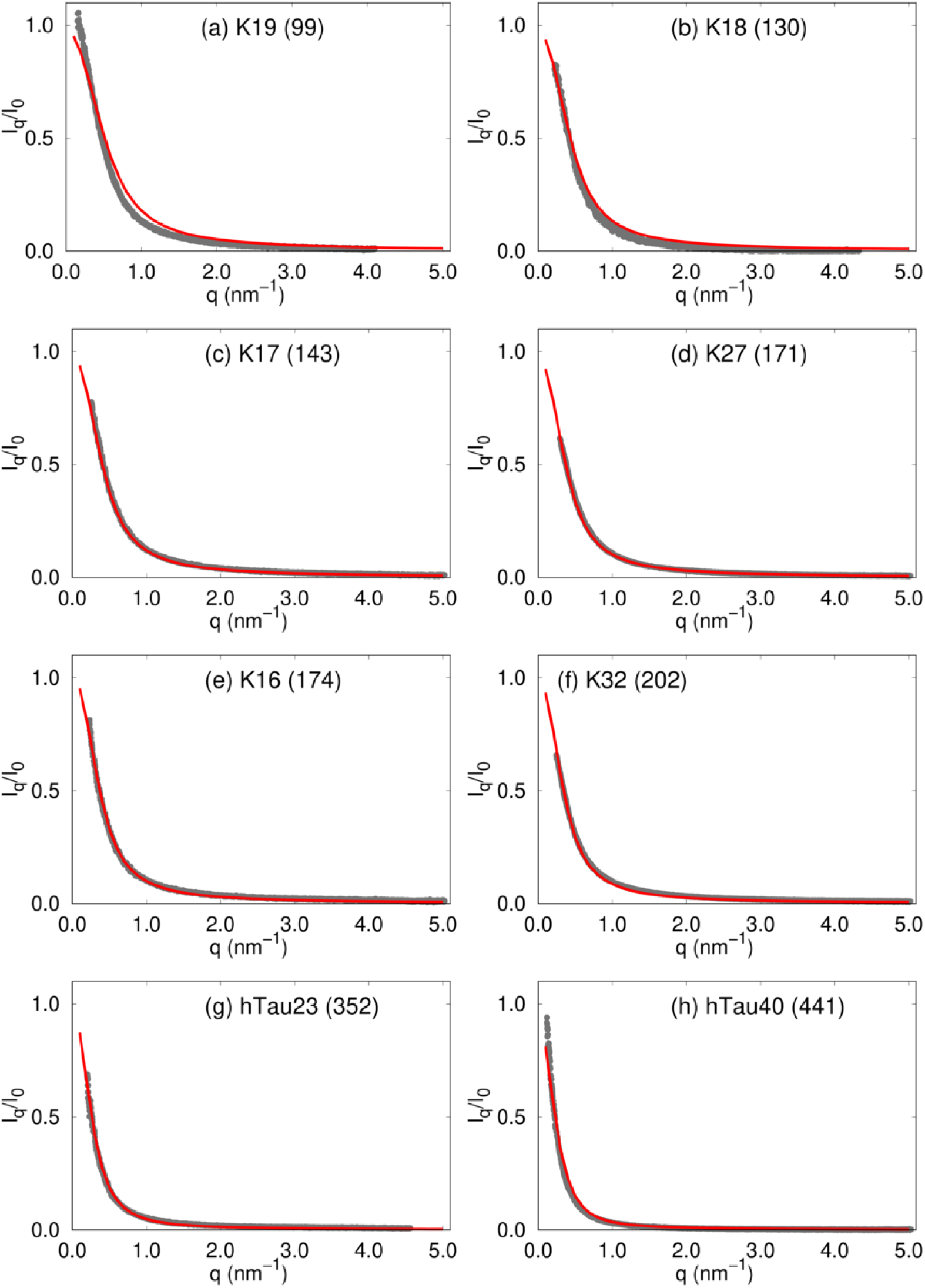
Comparison of SAXS profiles for eight Tau protein constructs. The lengths of the sequences are shown in parentheses in each figure. The color coding is the same as in Figure 1.

### Dependence of radius of gyration and hydrodynamic radius on sequence length

Given the accurate reproduction of the SAXS profiles (Figure 1 and Figure 2) for most IDPs, it is not surprising that the *R*_*g*_ values computed from simulations are also in accord with the experimental values (Figure S3). The scaling of *R*_*g*_ with sequence length (*N*_*T*_), fitted using the calculated or experimental *R*_*g*_ values, follows 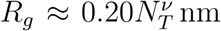, with *v* ≈ 0.588 (Figure 3 (a)), which naively suggests that in the experimental conditions the IDPs might be in a good solvent. However, this is not the case (see below).

**Figure 3:**
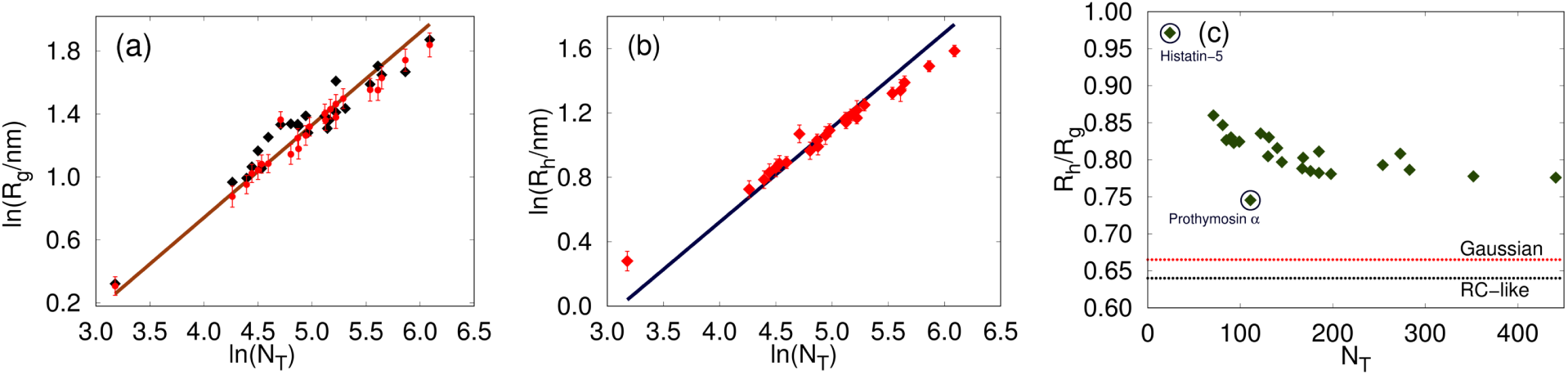
(a) Values of *R*_*g*_ from simulations (red circles) with comparison to the experimental estimates (black squares). The brown solid line is a fit to the power law: 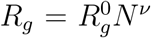, with 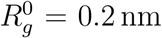 and *v* = 0.588. (b) Hydrodynamic radii, *R*_*h*_s, of the IDPs computed from the simulation trajectories (red symbols). The solid blue line is a power law fit: 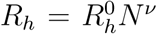, with 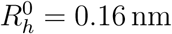 and *v* = 0.588. Both *R*_*g*_ and *R*_*h*_ follow the Flory random coil predictions. (c) The ratio of the hydrodynamic radius to the radius of gyration *R*_*h*_*/R*_*g*_, obtained from simulations. The outliers, Histatin-5 and Prothymosin-*α* are marked with circles. The theoretical limits are marked as dotted lines. Note that there are substantial deviations from the theoretical predictions for ideal chain and polymer in a good solvent.

We also calculated the hydrodynamic radii, *R*_*h*_, from the conformational ensemble using Eq. 5. The computed values of *R*_*h*_ are shown in Figure 3(b), and are tabulated in Table S3 of the SI. Though not equal in magnitude, both *R*_*g*_ and *R*_*h*_ (measurable from NMR for example) should show the same scaling with *N*_*T*_ with identical values of *v* (Figures 3 (a,b)).^59^ However, this scaling behavior does not imply that these IDPs behave as random coils.^32^ Indeed, the *R*_*h*_*/R*_*g*_ ratios from simulations deviate substantially from the established theoretical limits, which are 0.665 and 0.640, for an ideal chain, and a polymer in a good solvent, respectively (Figure 3(c)). As *N*_*T*_ increases beyond 300, *R*_*h*_*/R*_*g*_ becomes insensitive to sequence length (Figure 3(c)), indicating that the observed deviations may not be due to finite size effects.

### Distributions of radius of gyration (*R*_*g*_) and end-end distances (*R*_*ee*_)

If the solution conditions used in experiments are good solvents for IDPs, as the *R*_*g*_ and *R*_*h*_ scaling with *N*_*T*_ imply (Figure 3), then it should be reflected in the distribution *P* (*R*_*ee*_) of the the end-to-end distance (*R*_*ee*_), which is rigorously known for a polymer in a good solvent as well as an ideal chain. Therefore, comparing the simulated distributions to the rigorous functional forms provides a stringent test of the solvent quality. The *P* (*R*_*ee*_) distributions, with *R*_*ee*_ measured as the separation between the backbone atoms at the termini, are shown in Figures 4 (a,b) (see also supplementary Figures S12 and S13 in the SI). As a general rule, we find that the *P* (*R*_*ee*_) for IDPs closely resemble the theoretical result for a random coil. However, for specific sequences the distributions are more skewed towards the ideal or the Gaussian chain limit.^32,60^ The deviations from the random coil limit based on the theoretical predictions are likely due to non-local electrostatic interactions, and sequence composition (see below). Figure 4(a) shows that the *P* (*R*_*ee*_) distribution for Prothymosin-*α* differs substantially from both the theoretical limits. The deviation can be attributed to the high fraction of charged residues (∼ 58%).

**Figure 4:**
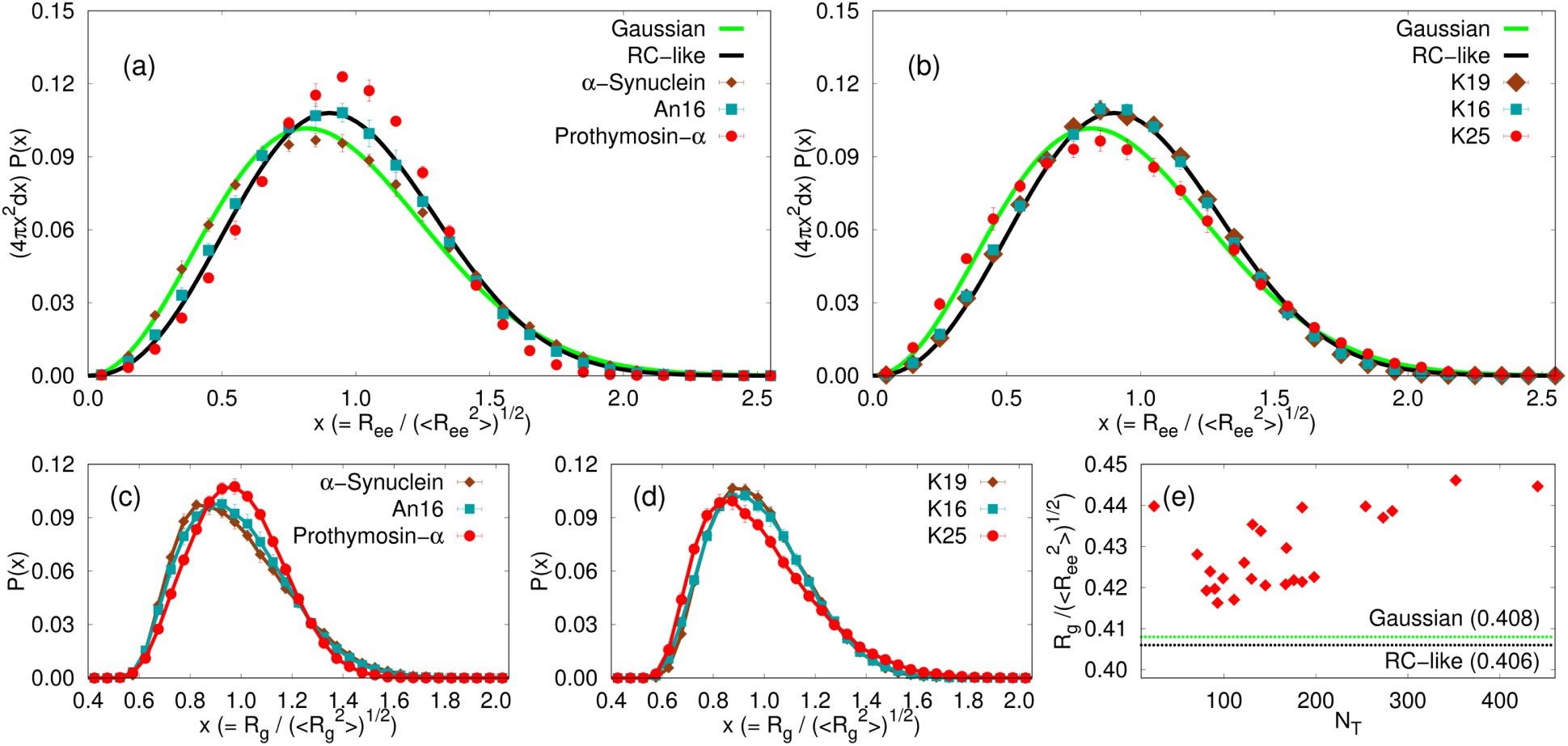
Distributions of the end to end distance (*R*_*ee*_), scaled by 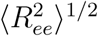. In order to compare with results for Gaussian chains (solid green line) and random coils (solid black line) we show 4*πx*^2^*P* (*x*), where 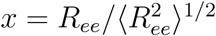. Figures (a) and (b) show the distributions for representatives from the sets of non-Tau IDP sequences and Tau-sequence constructs, respectively. If a particular IDP behaves strictly as a random coil (Gaussian chain) in terms of end-to-end distance distribution, then the corresponding distribution should coincide exactly with the theoretical result in black (green). (c,d) Distributions of *R*_*g*_ for IDPs described in (a) and (b) respectively. (e) Ratio of *R*_*g*_ to root mean square *R*_*ee*_, for all 24 IDPs described in the manuscript. The dashed horizontal lines in green and black show the ratio for a Gaussian chain and a random coil, respectively.

### Sequence matters

Interesting sequence-specific effects on the *P* (*R*_*ee*_)s are also observed for the Tau protein constructs, in spite of their common parent sequence. The *P* (*R*_*ee*_) for the majority of the Tau protein constructs show excellent agreement with the theoretical *P* (*R*_*ee*_) for a random coil (see Figure 4 (b) for K19 and K16, and also Figure S11 in the SI), and deviate substantially from that for an ideal chain. Deviations from the random coil behavior are found for 5 out of the 12 Tau sequence constructs, namely K25 (Figure 4 (b)), K23, K44, hTau23 and the WT sequence, hTau40 (Figure S13 in the SI). The *P* (*R*_*ee*_) for these sequences show increased propensities for smaller *R*_*ee*_ distances, as can be seen from Figure 4(b) for the K25 Tau construct. These sequence-specific deviations from average random coil-like behavior are also discernible in the distribution of *R*_*g*_ values, shown in Figures 4 (c) and (d). In Figure 4(e), the ratio *R*_*g*_ to root-mean-square *R*_*ee*_ is shown as a function of *N*_*T*_. This ratio is 0.406 for a random coil, and 0.408 for a Gaussian chain.^60^ Interestingly, even for those IDPs, whose *P* (*R*_*ee*_) coincides with the theoretical predictions for random coils, the 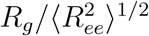 values deviate from the theoretical limits. This deviation becomes particularly pronounced as *N*_*T*_ increases, most notably for the longer Tau protein constructs.

The Tau sequences, which do not conform to standard polymer behavior, namely K25, K23, K44, hTau23, and hTau40, do not differ appreciably from the rest of the sequence constructs in terms of conventional sequence compositional parameters often used to rationalize their shapes (Table S4 in the SI). This apparent incongruity could be rationalized in terms of ensemble averaged contact maps obtained from simulations (see the Discussion section). We document below that the deviation from the theoretical random coil limit, is in fact a direct manifestation of the sequence-dependent conformational heterogeneity.

### Shape parameters are sequence dependent

The sequence-specific shape fluctuations of IDPs can be gleaned from the distributions of their shape parameters, Δ and *S* (see Eq. 6 in Methods). The calculated values of Δ, and *S* indicate that the conformations of IDPs can be described as prolate ellipsoids (Table S3 in the SI). In Figure 5, we show the distributions for four IDP sequences, and the rest are included in the supporting information (Figures S6–S9).

**Figure 5:**
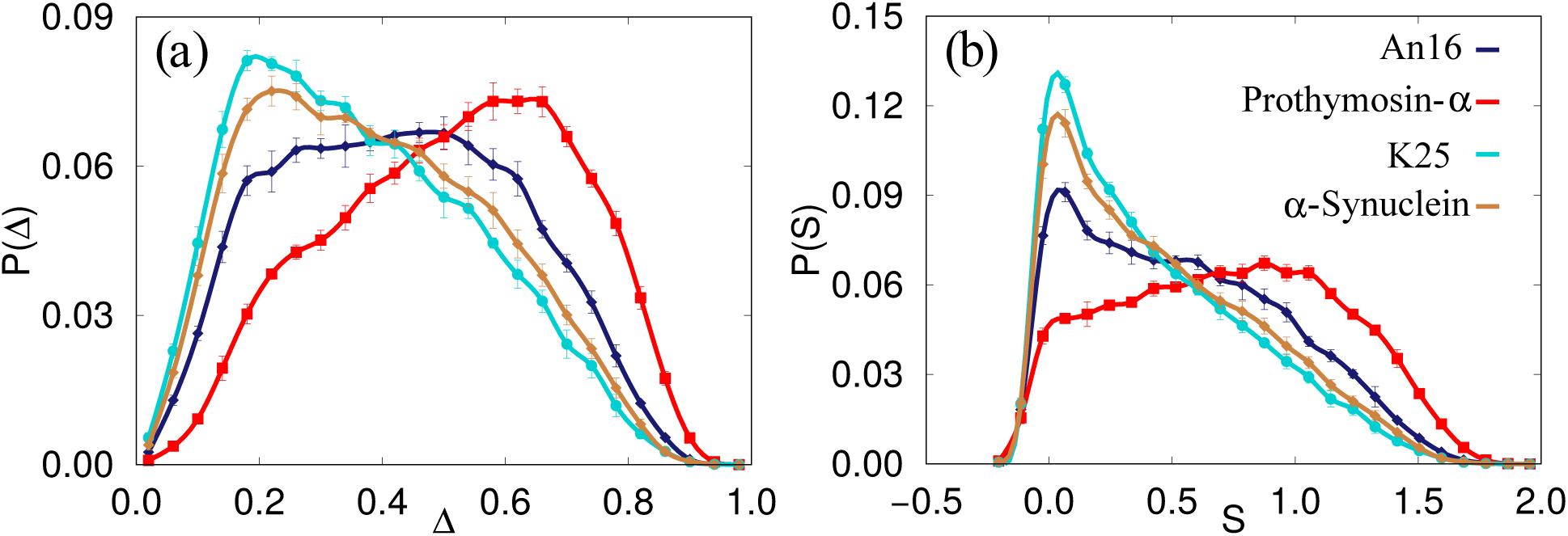
Comparison of the distributions of shape parameters (a) Δ and (b) *S* for An-16 (blue, diamonds) *α*-Synuclein (brown, diamonds), Prothymosin-*α* (red, squares) and K25 construct of the Tau protein (cyan, circles). Note that the average values of Δ and *S* for Gaussian chains are 0.52 and 0.87, and the corresponding estimates for an equivalent polymer in a good solvent are 0.55 and 0.91 respectively. The mean values of both Δ and *S* for the IDPs are drastically different from the expected theoretical values for standard polymer models.

Figure 5 (a) shows that Prothymosin-*α* has a high preference for extended conformations, while the structural ensembles for the K25 construct, and *α*-Synuclein are closer to being spherical. In contrast, the shape parameters for An-16 are homogeneously distributed, suggesting that elongated, and spherical conformations are equally probable. This systematic trend is also reflected in the distributions of *S* (Figure 5(b)), where the bias towards prolate conformations is maximal for Prothymosin-*α*, and minimal for K25. The large dispersions in *P* (Δ) and *P* (*S*) makes the calculation of the mean values, listed in Table S3 of the SI, not meaningful (see SI for elaboration). For comparison, the average values of Δ and *S* for Gaussian chains are 0.52 and 0.87, and the corresponding estimates for an equivalent polymer in a good solvent are 0.55 and 0.91 respectively.^61^ Overall, the shape fluctuations of the IDPs allude to a sequence-dependent heterogeneity of conformational ensembles. This aspect, which seems characteristic of the conformations of the IDPs explored here, is discussed in more detail below.

### IDP conformations are structurally heterogeneous

The distributions of *R*_*g*_ and *R*_*ee*_, as well as their deviations from random coil-like behavior for certain IDPs, suggest that the equilibrium populations of different conformations populating the IDP ensemble depend on the sequence. To illustrate the importance of conformational fluctuations in determining the statistical properties of IDPs, we consider representative examples: Prothymosin-*α, α*-Synuclein, An-16, and the K25 construct from the family of Tau proteins.

To obtain insights into the structural ensembles, and reveal their heterogeneous nature, we performed hierarchical clustering of the IDP conformations. Hierarchical clustering not only provides a means to quantify the contrasting features of the structural ensembles, which are evident from the Δ and *S* distributions, but also aids in visualizing the extent of the underlying conformational heterogeneity. We note that several authors have used ‘heterogeneity’ as a concept to underscore the differences between homopolymers and IDPs.^62–64^ More recently, this idea has also been suggested to explain the apparent lack of agreement between the *R*_*g*_ values measured using SAXS, and those inferred from FRET experiments.^65–67^

The results from the clustering analyses of *α*-Synuclein, and the K25 Tau construct are shown in Figure 6. The conformational ensembles corresponding to An16 and Prothymosin*α* are depicted in Figure 7. The appropriate number of clusters for each sequence was determined by evaluating the largest distance jumps in the corresponding dendrograms using the elbow method.^68^ Figures 6 and 7 show that our clustering scheme is robust, and the various families are clearly demarcated on a two-dimensional projection of the conformational landscape onto the *R*_*g*_, and *R*_*ee*_ coordinates.

**Figure 6:**
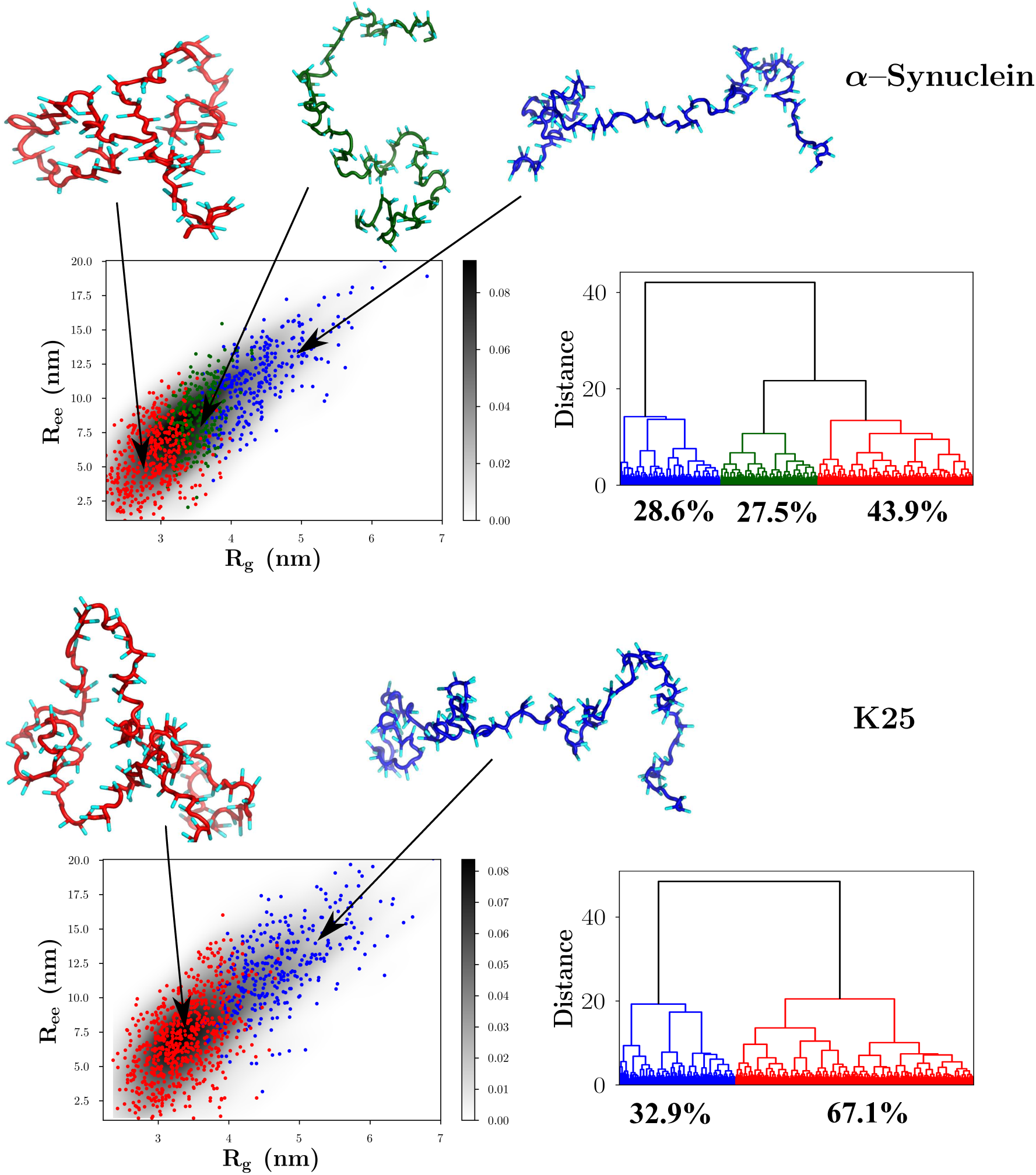
Hierarchical clustering of the IDP conformational ensembles using the Ward variance minimization algorithm. Upper panel: *α*-Synuclein; Lower panel: K25. The conformational landscapes (depicted as probability density plots) projected onto *R*_*g*_ and *R*_*ee*_ are shown on the left of each figure, and the corresponding dendrograms are shown on the right. Representative snapshots from each family, with their backbones rendered in different colors, are shown superposed on the conformational landscape. The same color-coding is used to classify clusters within the dendrograms, and the two-dimensional plots. The relative cluster populations are marked below the appropriate dendrogram branches.

**Figure 7:**
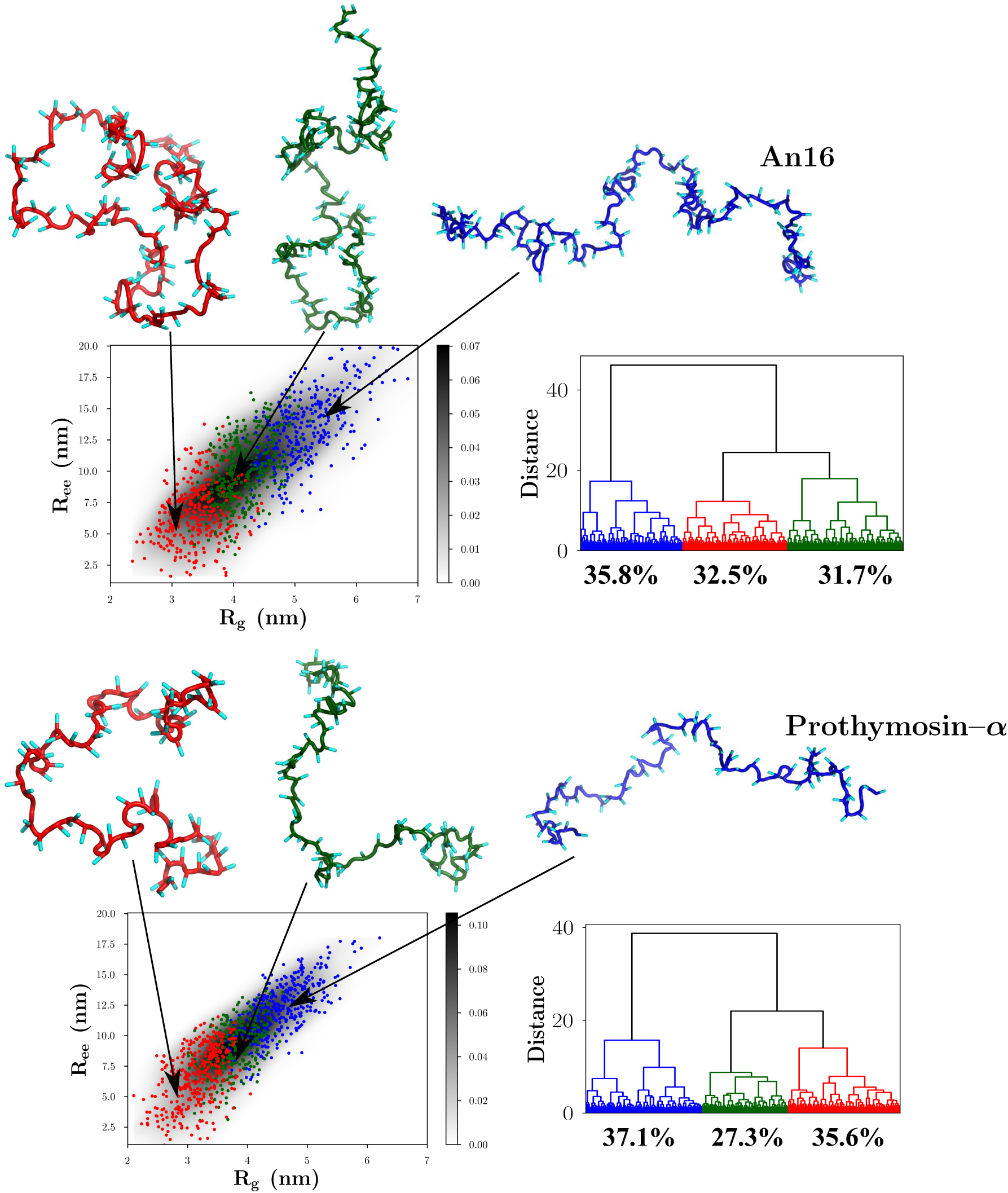
Upper panel: An16; Lower panel: Prothymosin-*α*. The conformational landscapes projected onto *R*_*g*_ and *R*_*ee*_ are shown on the left of each figure, and the corresponding dendrograms are shown on the right. Representative snapshots from each family, with their backbones rendered in different colors, are shown superposed on the conformational landscape. The same color-coding is used to classify clusters within the dendrograms, and the two-dimensional plots. The relative cluster populations are marked below the appropriate dendrogram branches.

#### α–Synuclein and K25

The equilibrium ensemble of *α*-Synuclein (Figure 6) partitions into at least three clusters, and consists of a large population of relative compact structures (43.9%). Both semi-extended, and fully extended structures are populated to a lesser extent, with occupation probabilities of 27.5%, and 28.6%, respectively. As is evident from the dendrogram (Figure 6), the equilibrium ensemble of the K25 Tau protein construct is overwhelmingly dominated by relatively compact structures, with the overall contribution being 67.1% to the net population. This trend is also reflected in the two-dimensional conformational landscape, where the region with small *R*_*g*_ and *R*_*ee*_ values is associated with the highest density. The shape of the conformational landscapes for both *α*-Synuclein, and K25, as well as the relative cluster sizes are commensurate with the observed deviations from random coil-like behavior.

#### An16 and Prothymosin-α

The equilibrium ensemble of An16 (Figure 7) consists of approximately equal contributions from relatively compact, semi-extended, and extended structures, with the populations being 32.5%, 31.7% and 35.8%, respectively. The relative populations suggest that no specific conformation is particularly favored for An16, which is consistent with a random coil-like behavior.

For the Prothymosin-*α* sequence, the ensemble of conformations partitions into at least three clusters (Figure 7). Although naively we expect that, due to the high value of net charge, there would be an overwhelmingly large population of extended structures for Prothymosin-*α*, clustering of conformations shows otherwise. While conformations exhibiting high *R*_*g*_ and *R*_*ee*_ values contribute maximally to the equilibrium population (37.1%), relatively compact conformations constitute the second-largest cluster, and account for as much as ≈ 35.6% of the conformational ensemble. Configurations having intermediate values of *R*_*g*_ and *R*_*ee*_, or the semi-extended structures have the lowest occupation probability (27.3%).

Overall, the systematic variations in the heterogeneity of sampled conformations for the different IDP sequences provide a structural explanation for the calculated distributions of *R*_*ee*_ and *R*_*g*_, which are clearly masked if only their mean values are analyzed. It is indeed remarkable that although mean values of *R*_*g*_ and *R*_*h*_ adhere to Flory scaling laws for good solvents, there is considerable fine structure in the IDP ensembles, as the clustering analyses demonstrate. It is likely that the plasticity of the conformational ensembles makes it possible for IDPs to carry out diverse functional roles in a bewildering array of metabolic pathways.

## Discussion

### Enhancements in local contact probability leads to deviation from random coil behavior

A hallmark of polymers in good solvents (random coils or RCs) is that the *P* (*R*_*ee*_) must obey the theoretical distribution obtained using polymer theory (see SI). The statistical properties of the ensemble of conformations of RCs are purely determined only by chain entropy. Interestingly, we find that although the dependence of *R*_*g*_ on *N*_*T*_ for all the IDPs is consistent with the Flory scaling law (Figure 3) there are deviations in *P* (*R*_*ee*_) from the expected theoretical predictions for RCs. For example, we observed such deviations for *α*-Synuclein, Prothymosin-*α*, as well as the Tau sequences K25, K23, K44, hTau23 and hTau40, but not for An16. The apparent conundrum can be understood from the dendrograms, which reveal clearly the sequence-dependent conformational heterogeneity. As a result the conformational ensembles are determined not only by entropy but also sequence-dependent energy, resulting in higher Boltzmann weight for certain classes of conformations relative to others.

To further illustrate the origin of deviation from the RC behavior, we calculated ensemble averaged contact maps from simulations using the SOP-IDP energy function, as well as the corresponding RC limit. We define the difference contact maps as,

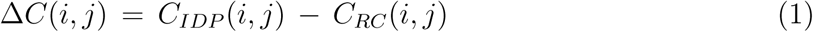

where *C*_*IDP*_ (*i, j*) is the contact probability between residues *i, j* of an IDP obtained from simulations, and *C*_*RC*_ (*i, j*) is the same contact probability obtained by specifically neglecting the contributions due to the last four terms in Eq. 2 (see the Methods section). Thus, Δ*C*(*i, j*) is a measure of the increased or decreased probability of contact between two residues, compared to an equivalent RC. Both *C*_*IDP*_ (*i, j*) and *C*_*RC*_ (*i, j*) were computed using the coordinates of the side-chain beads, including glycine and alanine. We assume that a contact exists if the distance between two residues is ≤ 0.8 nm.

The three contact maps shown in Figure 8 represent characteristically divergent scenarios.

**Figure 8:**
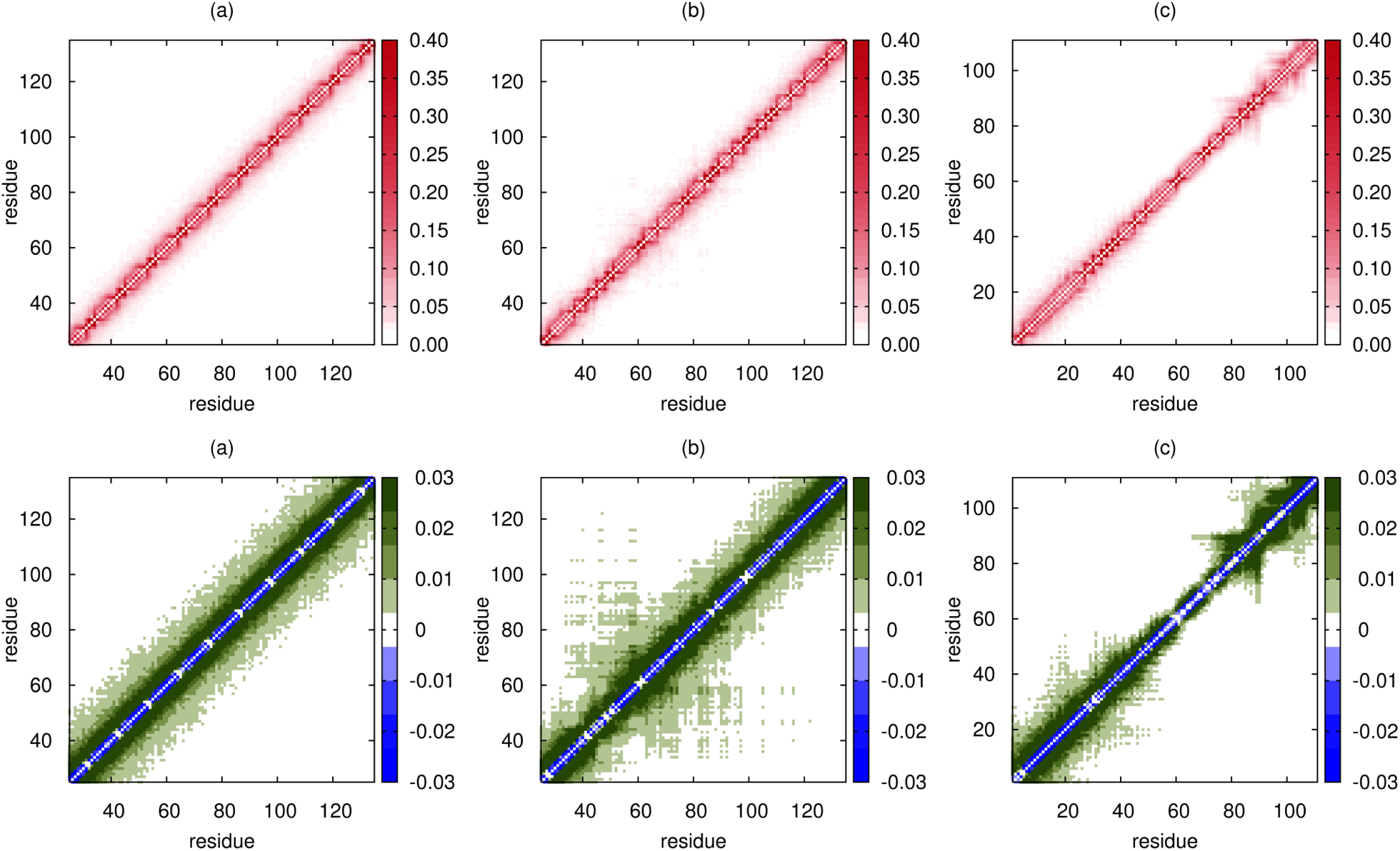
Contact maps for (a) An16, (b) the K25 Tau construct, and (c) Prothymosin-*α*. The top panel shows the contact maps for the three IDPs calculated using the SOP-IDP simulations. To illustrate the deviations for the corresponding RC contact map we plot the difference map using Eq. (1) (bottom panel). While the contact map for An16 reveals no sequence driven propensities for local structures, the latter two clearly highlight the presence of local domains that are more compact compared to the rest of the sequence. The nearly RC-like behavior of the mid-segment of Prothymosin-*α*, in stark contrast to the other IDPs studied (see also Figures S14-S16 in the SI), rationalizes the peripheral contribution of compact conformations observed for it in the hierarchical clustering of the conformational ensemble. Each of the contact maps depict a segment of 111 residues (*N*_*T*_ for Prothymosin-*α*) for optimal visual comparison. The full ranges for (a) and (b) are shown in Figure S16 in the SI.

#### An16

The contact map for An16 (Figure 8 (a)) shows a uniform enhancement in contacts between residues proximate along the sequence, over the entire sequence length compared to an equivalent RC polymer. This can readily be understood as the consequence of the non-specific attractive interactions between non-bonded residues, with sequence specific interactions (final term in Eq. 2) playing a relatively minor role. Without the formation of stable local structures, such enhanced contacts do not alter the RC-like nature of the polymer, as also evidenced by the RC-like *P* (*R*_*ee*_) distribution for An16 (Figure 4).

#### K25

The results in Figure 8 (b) show that the K25 Tau construct contains a locally compact segment, with preferentially enhanced contacts compared to the rest of the sequence. The segment with locally enhanced contacts is also present in the WT Tau sequence (hTau40), as well as the K25, K23, K44, and hTau23 constructs (shown in Figures S15 and S16). We infer that the presence of this region, which results in the formation of the locally preferred structures, leads to the observed deviations from the RC-like behavior for the five sequence constructs.

#### Prothymosin-α

In this IDP (Figure 8 (c)), the two termini adopt locally compact structures, while the mid-segment of the IDP closely conforms to the RC-limit. For this IDP, electrostatic interactions dominate the conformational properties, and the termini, with both positive- and negatively charged residues, undergo local compaction. However, both the termini are overall negative, leading to unfavorable local electrostatic interactions. Both these effects together lead to the predominance of extended, as well as relatively conformations in the conformational ensemble (Figure 7).

### Structural ensemble of *α*-Synuclein

Our simulations provide considerable insights into the structural details of *α*-Synuclein, which has been extensively investigated using a variety of experimental methods. The calculated hydrodynamic radius reported in Table S3 (*R*_*h*_ = 2.89 *±* 0.17) nm is in very good agreement with experimental value (*R*_*h*_ = 2.74 *±* 0.04) nm obtained using pulse field gradient NMR.^69^ Similarly, the calculated *R*_*g*_ value is in close agreement with experimental estimate (see Table S3), with the difference between simulation and experimental estimates being about 12%. More importantly, our simulations provide a complete characterization of the subtle structural details that are difficult to glean from experiments. First, the segregation of the conformational landscape for *α*-Synuclein is in accord with recent experimental works^70–73^ that hinted at the presence of distinct conformational states in the structural ensemble, with compact structures having the highest population at equilibrium. Specifically, the difference contact map (Figure 9) reveals that there are long range tertiary interactions between residues in the acidic C-terminal region, and the NAC domain in *α*-Synuclein. Previous works^74,75^ suggest that such interactions, which protect the NAC region against fibrillation, play a key role in impeding aggregation. Interestingly, tertiary contacts between residues in the neighborhood of 74 and 94, first noted in experiments using electron transfer,^76,77^ are captured in the contact maps presented in Figure 9. Because of large conformational fluctuations such tertiary interactions, which could serve as initiating nucleation sites for aggregation, the probability of forming such contacts is not substantial but detectable.

**Figure 9:**
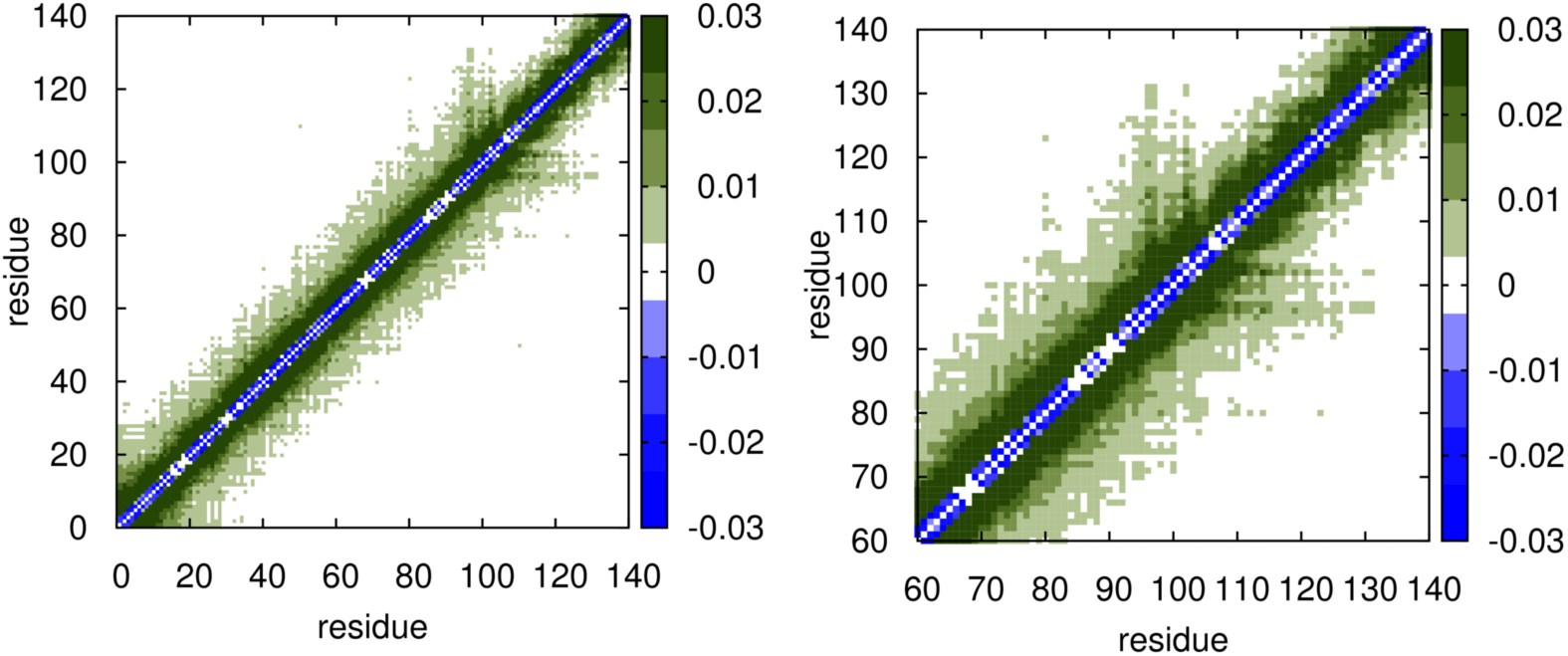
Difference contact maps for the *α*-Synuclein. Left: whole sequence Right: zoomed-in view highlighting the contacts between the C-terminal and the NAC regions.

Sequence-specific deviations from the RC-limit highlight the heterogeneity in the population of IDPs. We reiterate that it cannot be trivially ascertained that water (or solution conditions used in experiments) act uniformly as a good solvent over the entire peptide sequence of an IDP. To classify IDP sequences as polymers in a good solvent, based solely on estimates of the Flory exponent *ν* with modest variations in *N*_*T*_ is erroneous, which severely undermines the heterogeneity of their conformations.

### Limitations of the SOP-IDP model

Although the SOP-IDP model is remarkably successful in capturing many aspects of the structural ensembles of even large IDPs, like all other empirical potentials, it is not without limitations. The SOP-IDP force field does reproduce quantitatively the radius of gyration of globular proteins at high denaturant concentrations, thus showing that the model is accurate in describing statistical properties of disordered states. It is unlikely that the model could predict the folding properties of even small globular proteins (for example the heat capacity) as a function of denaturant concentration as accurately as we have done here for *I*(*q*) for IDPs.

Recently, the goal^35,37,38^ of creating a transferable force field, which yields highly accurate results for both globular proteins and IDPs from essentially sequence alone, has been a motivating factor to drastically alter the currently popular force fields. Although laudable, achieving this goal is a daunting task because it is tantamount to solving all aspects of the protein folding problem and issues in IDPs in one fell swoop. In these studies,,^35,37,38^ which use three entirely different empirical energy functions, the calculated *I*(*q*)s for the 24-residue RS peptide (see the SI for details) and the 71-residue ACTR IDPs are in very good agreement with experiments. In addition, the *R*_*g*_ values for a few IDPs calculated using a force field that differs greatly from the ones used by others^35,38^ agree with measurements. However, for foldable peptides and miniproteins the agreement between simulations and experiments is not satisfactory, which means the stated goal of creating a transferable force field has not been accomplished. This illustrates the difficulty in building a truly “universal” potential within the limitations inherent in additive classical force fields, as articulated elsewhere.^38^ With the possible exception of a recent study of IDPs,^37^ applications (especially to globular proteins) are restricted to small *N*_*T*_ values, thus leaving open the question of the extent of transferability for longer *N*_*T*_ even when restricted to IDPs. It is indeed the case that a large number of energy functions, at all levels of coarse-graining, could be constructed for successful applications to possibly only a certain class of proteins.

The SOP-IDP force field is not designed to be a universal potential that can simultaneously describe accurately the measurable properties of both IDPs and globular proteins. We have shown that the SOP-IDP force field is unique in accuracy for IDPs, providing a new perspective for describing sequence-specific properties of intrinsically disordered proteins (and by statistical equivalence the DSE of globular proteins at high denaturant concentrations). However, for structural characterization of folded proteins under physiological conditions, as well as studies of various aspects of protein folding, the parameters described in Liu *et al.*^78^ are appropriate. The one advantage is that the SOP model has only three independent parameters, and hence a broad range of applications could be explored to assess its limitations.

### Additional tests

We have taken advantage of the availability of experimental data and results from atomic detailed simulations of Wu *et al.*^38^ to carry out simulations for a 24-residue RS peptide using the SOP-IDP model. Wu *et al.*^38^ used the RSFF2+/TIP4P-D force field to calculate the scattering profile (*I*(*q*) vs *q*) for the RS peptide, and found it to be in excellent agreement with SAXS experiments. We used the current model to simulate the RS peptide, and find that the *I*(*q*)s from the SOP-IDP model, the all-atom simulations and the measured SAXS profile nicely overlap with each other (see Fig. S4 in the SI). In the SI, we also illustrate additional applications to the 20-residue GS10 and the 26-residue HIV-1 Rev peptides for which *R*_*g*_ values (but not SAXS profiles) have been reported by Wu *et al.*^38^ The results using coarse-grained and RSFF2+/TIP4P-D force field are in good agreement. It should be pointed out that a similar level level of accuracy could be obtained using the Flory formula given in the caption to Figure 3a, without the need for *any* simulations, which shows that success of a force field can only be assessed by comparing the simulated and measured SAXS profiles. The predictions in the SI as well the results in the main text show vividly that the proposed SOP-IDP energy function is highly efficacious in obtaining statistical properties of 27 IDPs of which 5 were used in the training set.

## Conclusions

Given the multifarious roles that IDPs play in a variety of cellular functions, there is an urgent need to decipher the physical principles governing IDP structure, and dynamics, at the molecular level. We have introduced a robust coarse-grained model for IDPs (SOP-IDP), which quantitatvely reproduces the SAXS profiles for a diverse range of sequences, and length varying from 24 to 441 residues. The parameters of the force field were calibrated via a learning procedure, using a few test sequences, and once optimized they were unaltered for 24 other IDP sequences of varying composition, charge densities, and sequence lengths.

Although globally the sizes of the IDPs follow the well known Flory scaling law obtained for synthetic polymers, equilibrium populations of different substates strongly depend on the precise IDP sequence. In the conformational substates of IDPs, we find evidence for specific weak intramolecular interactions between certain residues, giving rise to population of conformations with local structures. We also show that the adaptability of IDPs arising from their conformational plasticity, which cannot be anticipated using sequence characteristics (such as fraction of positive and negatively charged residues) alone,^27^ is important in their ability to interact with a multitude of partners to form fuzzy complexes stabilized by specific intermolecular interactions. Our work lends further credence to an existing viewpoint that the observed sequence-dependent conformational heterogeneity determines the functions of IDPs. The accuracy of the computations in describing the equilibrium ensemble of a number of IDPs sets the stage for describing IDP-RNA interactions, which is needed to understand ribonucleoprotein stress granule formation in eukaryotes, as well as quantitative descriptions of the formation of biomolecular condensates at the molecular level.

## Methods

### Development of the Model

In the Self-Organized Polymer (SOP-IDP) model, except for glycine and alanine, the amino acid residues are represented using a backbone and a side chain (SC) bead. We use the *C*_*α*_ atoms to model glycine and alanine owing to their small side chains. The radii of the individual beads are given in Table S1 of the SI. The energy function in the SOP-IDP model is:

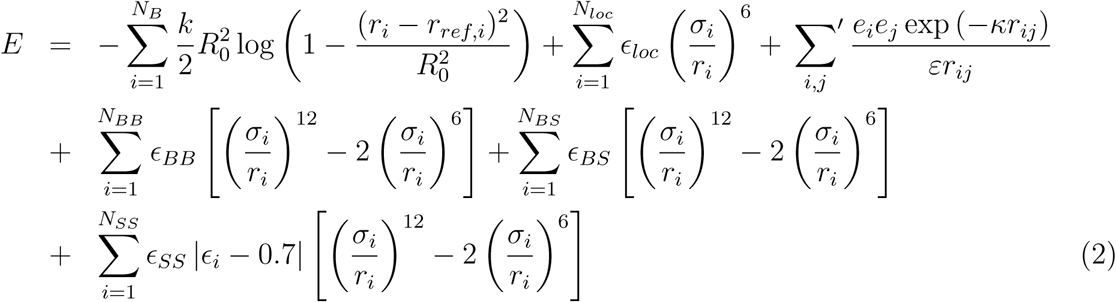

The first term in Eq. 2, for a chain with *N*_*B*_ bonds, represents bonded interactions, which are described using a finite non-linear elastic (FENE) potential. The purely repulsive second term acts only between bead pairs (total number *N*_*loc*_) that are not covalently bonded, but belong to residues separated by ≤ 2 along the polypeptide chain. It represents the excluded volume interactions that prevent unphysical bond-crossing between pairs of beads that closely follow one another along the polymer chain. The third term is a screened Coulomb potential, that accounts for electrostatic interactions between all pairs of charged residues. The charge corresponding to a residue is assigned to the SC. The titratable residue histidine is treated as neutral in the model, unless it is experimentally determined to be in the protonated state. The parameters *κ* and *ε* in the third term in Eq. 2 are the inverse Debye length, and the dielectric constant, respectively. These three terms were previously used in protein folding and related studies,^43,44,79^ where the justifications for their choices are given.

The final three terms in Eq. 2 represent interactions between backbone–backbone (BB), backbone–side chain (BS), and side chain–side chain (SS) beads, respectively. The total numbers of such BB, BS, and SS pairs, are respectively, *N*_*BB*_, *N*_*BS*_ and *N*_*SS*_. Interactions between the side chains depend only on the identities of the amino acids without bias towards any specific structure. The parameter *ϵ*_*i*_ in the final term in Eq. 2 is obtained from the Betancourt-Thirumalai statistical potential.^80^ The *ϵ*_*i*_ values are unique to each pair of amino acid residues, and hence, depend explicitly on the IDP sequence. In Eq. 2, the three parameters *ϵ*_*BB*_, *ϵ*_*BS*_ and *ϵ*_*SS*_ set the energy scales corresponding to the non-local interactions, and are the only free parameters in the energy function. Following the standard convention used in polymer physics, these three parameters are expressed in units of *k*_*B*_*T* (with T=298 K). We determine the initial values of *ϵ*_*BB*_, *ϵ*_*BS*_ and *ϵ*_*SS*_ in a top-down fashion, by using SAXS data (*R*_*g*_, and *I*(*q*))for only three short IDPs (24 ≤ *N*_*T*_ ≤ 131) as constraints in the parametrization scheme. Subsequently, to find the optimum values of the parameters, only the experimental *R*_*g*_ estimates for three long IDP sequences (202 ≤ *N*_*T*_ ≤ 441) were used as constraints in the parametrization scheme. Further details of this learning procedure are provided in the supporting information.

### Comments on the parameter values

The values of the parameters, which were obtained using a learning procedure described in the SI, in the SOP-IDP model are (in units of *k*_*B*_*T*):

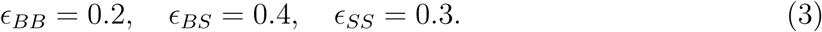

These values differ from the ones used to describe globular proteins,^78^ which were obtained in order to describe their folding thermodynamics at zero denaturant concentrations ([C]s). As [C] increases, the changes in the stability of the folded states decreases, which we accounted for phenomenologically using transfer free energies, thus creating the SOP-MTM model.^78^ The effective interaction parameters (*ϵ*_*BB*_, *ϵ*_*BS*_, *ϵ*_*SS*_) are expected to decrease as [C] increases, so that at high [C], the statistical properties of the denatured state ensemble (DSE) of globular proteins and IDPs, such as *R*_*g*_, would exhibit the scaling 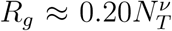, with *v* ≈ 0.588.^81^ Because the *R*_*g*_ scaling of DSEs of globular proteins at high denaturant concentrations is statistically equivalent to those for IDPs (hence the plausible relevance of IDPs to the SAXS-FRET controversy), it is not surprising that the optimal values for the SOP-IDP model (Eq. 3) differ from the [C] = 0 values.^78^ The values of *ϵ*_*BB*_, *ϵ*_*BS*_, and *ϵ*_*SS*_ for globular proteins in water, respectively, are 4.6, 1.7 and 1.7 times larger than the values for IDPs (Eq. 3).

If the reasoning above is correct then we ought to obtain reasonably accurate *R*_*g*_ values for globular proteins at high denaturant concentrations using the parameters given in Eq. 3. After all, at high [C], unfolded states of globular proteins are statistically equivalent to IDPs. To this end, we simulated the sequence corresponding to to the highly charged Ubiquitin (PDB ID: 1UBQ) using the current SOP-IDP force field. The estimated *R*_*g*_ for Ubiquitin using the SOP-IDP model is 2.51 and 2.55 nm at 150 and 1000 mM monovalent salt concentrations, respectively. These values compare well with the *R*_*g*_ for the DSE of Ubiquitin (≈ 2.56 nm, see for example, Fig 4A in Gates *et al.*,^81^ and Fig. 2A in Reddy *et al.*^82^ at high denaturant concentration). The simulations described in Reddy *et al* ^82^ were carried out using the parameters described in Liu *et al.*,^78^ but taking into account the effect of denaturants using MTM. These new simulations for Ubiquitin using the SOP-IDP model are gratifying because the SOP-MTM,^82^ and the current model were developed using entirely different methods for different purposes. This shows that the parameters for the IDP model describe well the properties of globular proteins at high denaturant concentrations (8 M GdmCl), in line with the expected statistical equivalence of their ensembles with IDPs. However, as pointed out in the Discussion section, the parameters will not be accurate in the sense found for IDPs, for real globular proteins.

### Simulations

We used the LAMMPS molecular dynamics simulator for all the simulations,^83^ which were performed using the under-damped Langevin dynamics.^84^ The equations of motion were integrated with a time step of 30 fs. To obtain statistically meaningful results, we carried out ten independent simulations for each system, which we ascertained are sufficient to obtain converged results. Each trajectory was equilibrated for 10^8^ simulation steps, following which the production runs were carried out for 2 *×* 10^8^ simulation steps for sequence lengths *<* 130, and 5 *×* 10^8^ simulation steps for longer sequences. Visual molecular dynamics (VMD) was used for visualization, as well as some data analyses.^85^

In our simulations, all the non-bonded interaction terms in Eq. 2 were implemented in their truncated and shifted forms, with the cut-off separation for short-range interactions set at 2.4 nm. The cut-off separation for the screened Coulomb potential was consistently chosen to be greater than 4*κ*^−1^. The inverse Debye length (*κ*) was determined from the corresponding monovalent salt concentrations employed in the simulations. We set the dielectric constant, *ε* = 78.

### Data Analyses

The simulated SAXS profiles were computed using the Debye formula,

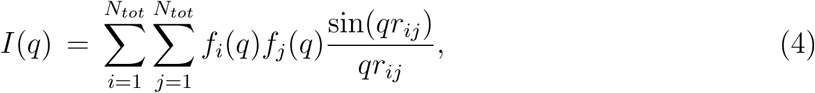

with *q*-dependent structure factors (*f* (*q*)), which were reported elsewhere.^86^ In Eq. 4, *N*_*tot*_ is the total number of beads in a given IDP.

### Hydrodynamic radius, *R*_*h*_

Dynamic light scattering (DLS), and fluorescence correlation spectroscopy (FCS) experiments^9,87–89^ are routinely used to measure *R*_*h*_. The value for *R*_*h*_ for a polymer is calculated as the radius of a hard sphere that has the same effective diffusive behavior as the polymer. We computed *R*_*h*_ from the simulations using:

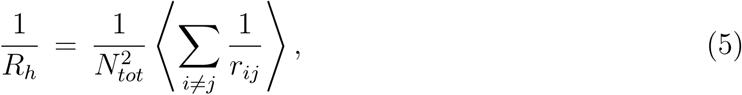

where *N*_*tot*_ is the total number of beads in the IDP model, and *r*_*ij*_ is the distance between beads *j* and *i*. The angular braces denote the ensemble average.

### Shape Parameters

We use Δ and *S* to characterize the shapes of folded and unfolded proteins,^90,91^ which are defined as,

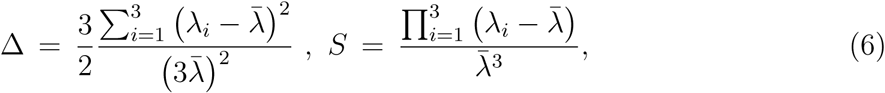

where *λ*_*i*_ are the eigenvalues of the gyration tensor, and 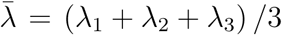.^90^ By definition, 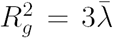. The Δ parameter characterizes the asphericity of conformations, and is bound between 0 (perfectly spherical) and 1 (linear). The shape parameter *S*, which is negative (positive) for oblate (prolate) ellipsoids satisfies the bound 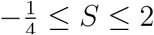.^92^

### Hierarchical Clustering

To identify representative conformations populating the IDP ensembles, as well as characterize their structural heterogeneity, we performed hierarchical clustering^93^ using a pairwise distance metric *D*_*ij*_, defined as,

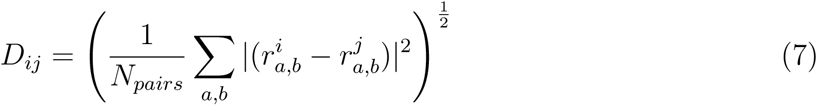

where 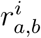 and 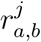 are the pairwise distances between the *C*_*α*_ atoms *a* and *b*, in snapshots *i* and *j*, respectively; *N*_*pairs*_ is the total number of *C*_*α*_ pairs. The terminal *C*_*α*_ atoms were excluded in the evaluation of the distance matrix. The Ward variance minimization criterion,^94^ as available within the *scipy* module, was employed to identify the distinct clusters. The hierarchical organization of clusters was visualized in the form of dendrograms. The structure exhibiting the lowest root-mean-square-deviation (RMSD) with respect to all the other members is identified as the representative structure of a given cluster.

## Acknowledgement

We are indebted to Prof. D. I. Svergun for providing us with the tabulated forms of the scattering profiles for the Tau sequences, which enabled us to calibrate the SOP-IDP model. We thank Robert Best for providing the latest unpublished SAXS data for ACTR, and Liz Rhoades for pointing out pertinent references, and Monika Fuxreiter for useful discussions. The research was supported by the National Science Foundation (CHE 16-36424), the Collie Welch Regents Chair (F-0019), and the National Institutes of Health (R01 GM107703 and GM089685). We are grateful to the Texas Advanced Computing Center (TACC) for providing the needed computing time.

## Supporting Information Available

Details of the learning procedure used to calibrate the SOP-IDP model; Distributions of the shape parameters for all the 24 IDP sequences; End-to-end (*R*_*ee*_) distributions for all the 24 IDP sequences; Sequence compositional properties of the IDP sequences; Contact maps for selected IDP sequences; FASTA representation of the IDP sequences.

This material is available free of charge via the Internet at http://pubs.acs.org/.

## Electronic supplementary information

### Development of the SOP-IDP model

We set ourself the task of creating a minimal model, which could be used for IDPs with arbitrary length and sequence, in order to accurately describe not only the average properties of IDPs but also the details of the structural ensemble. Although not investigated here, the model ought to be simple enough to include the effects of denaturants, as is often done in experiments. To this end, we created the SOP-IDP model, which is built on the successful Self-Organized Polymer (SOP) model used to study temperature and denaturant dependent folding thermodynamics and kinetics of a large number of globular proteins.^1,2^ Unlike in the case of globular protein folding, where neglecting non-native interactions may be justified,^3,4^ in describing IDPs all amino residues have to be treated on equal footing. In the SOP-IDP model each residue is represented by a C_*α*_ atom, and a side chain (SC) bead that is covalently bonded to the C_*α*_ atom. Exceptions to this representation are glycine and alanine, which are represented in the SOP-IDP model by single beads owing to their small sizes. In the implementation of the pair potentials, the glycine and alanine beads are thus treated as both side chain beads, and backbone beads, depending on the type of the partner. The interactions between pairs of glycine or alanine beads are treated through the SC-SC interaction potential in the energy function, to account for sequence specificity. The charges and the van der Waals radii, which are needed for integrating the low friction equations of motion using the SOP-IDP force field, for all the interaction sites are given in Table S1.

**Table S1:**
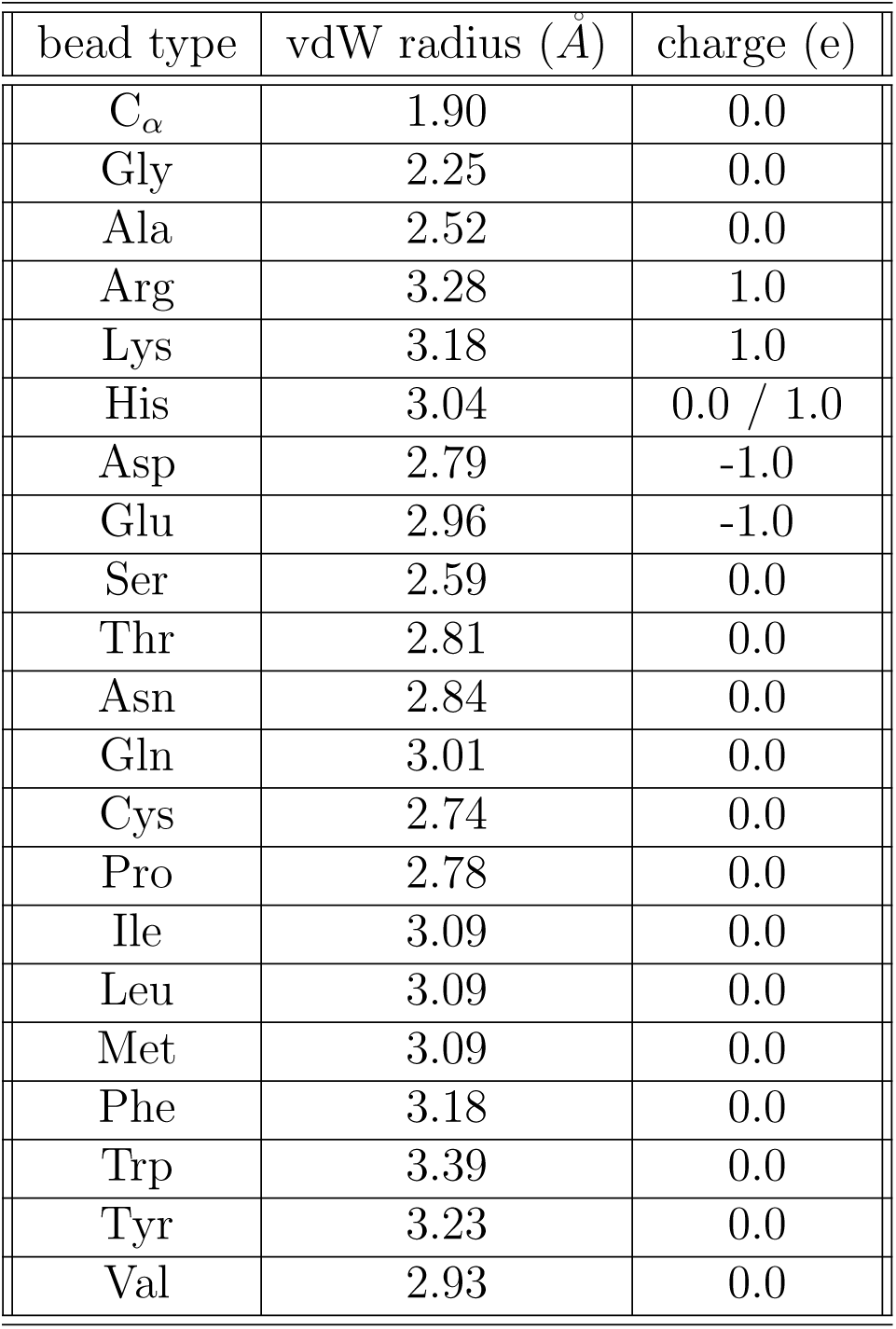
Parameters for the coarse-grained beads in the SOP-IDP model.

### Learning procedure for parametrizing the SOP-IDP model

The pre-factors *ϵ*_*BB*_, *ϵ*_*BS*_, and *ϵ*_*SS*_, which set the energy scales corresponding to the nonlocal interactions (see Eq. 1 in the main text), are the only free parameters in the SOP-IDP energy function. We used the experimental estimates of *R*_*g*_ as well as the low *q* regions of the SAXS profiles for three relatively short IDP sequences, Histatin-5, ACTR, and hNHE1, to obtain the initial estimates for the three free parameters, *ϵ*_*BB*_, *ϵ*_*BS*_, *ϵ*_*SS*_ in the SOP-IDP model.^5,6^ Histatin-5 is a small (24 residue) IDP that has often been used as reference for testing the validity of computational models.^5,7^ The other two IDPs, ACTR (*N*_*T*_ = 71) and hNHE1 (*N*_*T*_ = 131) are also well studied.^6,8^ In addition, the SAXS profiles for Histatin-5, ACTR and hNHE1 at 150 mM monovalent salt concentration have low noise-to-signal ratio, which makes objective comparisons feasible with the simulated profiles at equivalent ionic strengths.

However, Histatin-5, ACTR, and hNHE1 are short compared to other frequently studied IDP sequences. Thus, we expanded our training set, and we refined the SOP-IDP energy function by using experimental *R*_*g*_ values for the K32, K23, and hTau40 sequences. The optimal set of parameters for the SOP-IDP model from our learning procedure is (in units of *k*_*B*_*T*):

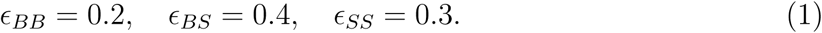

The values quoted in Eq. 1 are different from the ones used for globular proteins.^1^ The maximum decrease is in *ϵ*_*BB*_, which is 4.6 times smaller than used previously whereas *ϵ*_*BS*_ and *ϵ*_*BS*_ are only ≈ 1.7 times smaller (see Table S1 in Liu *et al.*^1^). Because the largest change is in *ϵ*_*BB*_, we recalculated *R*_*g*_ for Histatin-5 using the value for *ϵ*_*BB*_ used previously.^1^ The resulting value is only about 16% smaller than the what is reported in Table S3.

Our rationale for seeking a different set of parameters are summarized below. (1) the values for globular proteins describe the situation with no denaturants (concentration of denaturants [*C*] = 0) so that the folded states are predominantly populated. As [*C*] increases, the stabilities of the folded states decrease, which we accounted for phenomenologically using transfer free energies, thus creating the SOP-MTM model.^9,10^ Within this framework, the relevant interaction energy scales would decrease linearly. For example, *ϵ*_*BS*_ *≈ ϵ*_*BS*_([*C*] = 0) − *m*_*BS*_[*C*] where *m*_*BS*_ is the analogue of the *m* value accounting for loss of global stability at non-zero [*C*]. (2) Because the ensemble of conformations of IDP behave like the unfolded states of globular proteins, created at high denaturant concentrations we reasoned that it is not appropriate to use the values that stabilize the folded states.

The current SOP-IDP model describes IDPs and denatured states of globular proteins. It cannot describe with near quantitative accuracy the states of globular proteins at all values of denaturant concentrations. We hasten to add that using the SOP-IDP parameters in Eq.1, we find that the mean *R*_*g*_ of the hairpin from GB1 protein is ≈ 1.04 nm, which compares favorably with the value (*R*_*g*_ ≈ 1.22 nm) calculated using the PDB structure. However, we should emphasize that there is no guarantee that the SOP-IDP model could describe the fate of globular proteins of arbitrary size and topology, at various external conditions, as accurately as we have done here for IDPs. Indeed, currently, no such force field at any level of description exists that can achieve this goal, and it is unlikely that a universal force field could be constructed, which would be accurate (errors in directly measurable quantities, when compared to experiments, that are small for a number of systems over a range of external conditions) for both globular proteins and IDPs. Construction of such a force field would be equivalent to solving the protein folding problem (prediction of structure, thermodynamics, and kinetics) from sequence alone.

### Quantitative comparison between the simulated and experimental SAXS profiles

The level of agreement between the simulated and experimental SAXS profiles (in the Kratky representation) for the 24 IDP sequences (whose scattering profiles are depicted in Fig. 1 and Fig. 2 in the main text) could be quantified by calculating the extent of deviation between the simulated and experimental SAXS profiles. From the simulation trajectories, the *q*^2^*I*_*q*_*/I*_0_ values were computed for each IDP sequence at discrete intervals of *q* (Δ*q* = 0.01*Å-*1). For a quantitative comparison, the number of data points available from the experimental SAXS profiles were reduced to produce datasets equivalent to those obtained from simulations.

Following the discretization, we calculated,

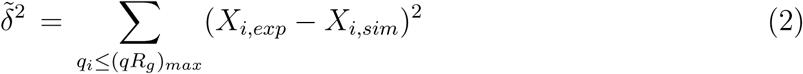

where 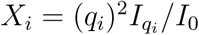 and [0,(*qR*_*g*_)_*max*_] is the range over which 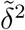 is estimated. In order to compare the relative errors for all the IDPs on equal footing, we report the error estimates in terms of *δ*^2^, which is defined as:

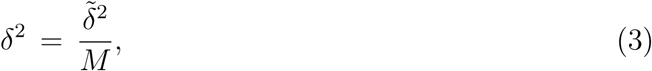

*M* being the number of data points available for comparison between the experimental and simulated data for a given IDP and for a given choice of (*qR*_*g*_)_*max*_.

The *δ*^2^ estimates corresponding to the simulated SAXS profiles (shown in Figs 1 and 2 in the main text) are tabulated in Table S2 for two different choices of (*qR*_*g*_)_*max*_. Table S2 shows that for values of (*qR*_*g*_)_*max*_ up to 3 (well beyond the normal Guinier regime) the relative errors are small while they increase at large (*qR*_*g*_)_*max*_. The results in Table S2 show that the predictions of the SOP-IDP model are fairly accurate.

### Comparison between simulated and experimental *R*_*g*_

In contrast to SAXS experiments, where *R*_*g*_ values are usually determined from a Guinier analysis of the scattering profiles in the low *q* regime, we can obtain *R*_*g*_ directly from simulations, using the standard polymer physics formula:

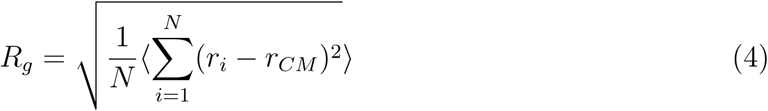

**Figure S1:**
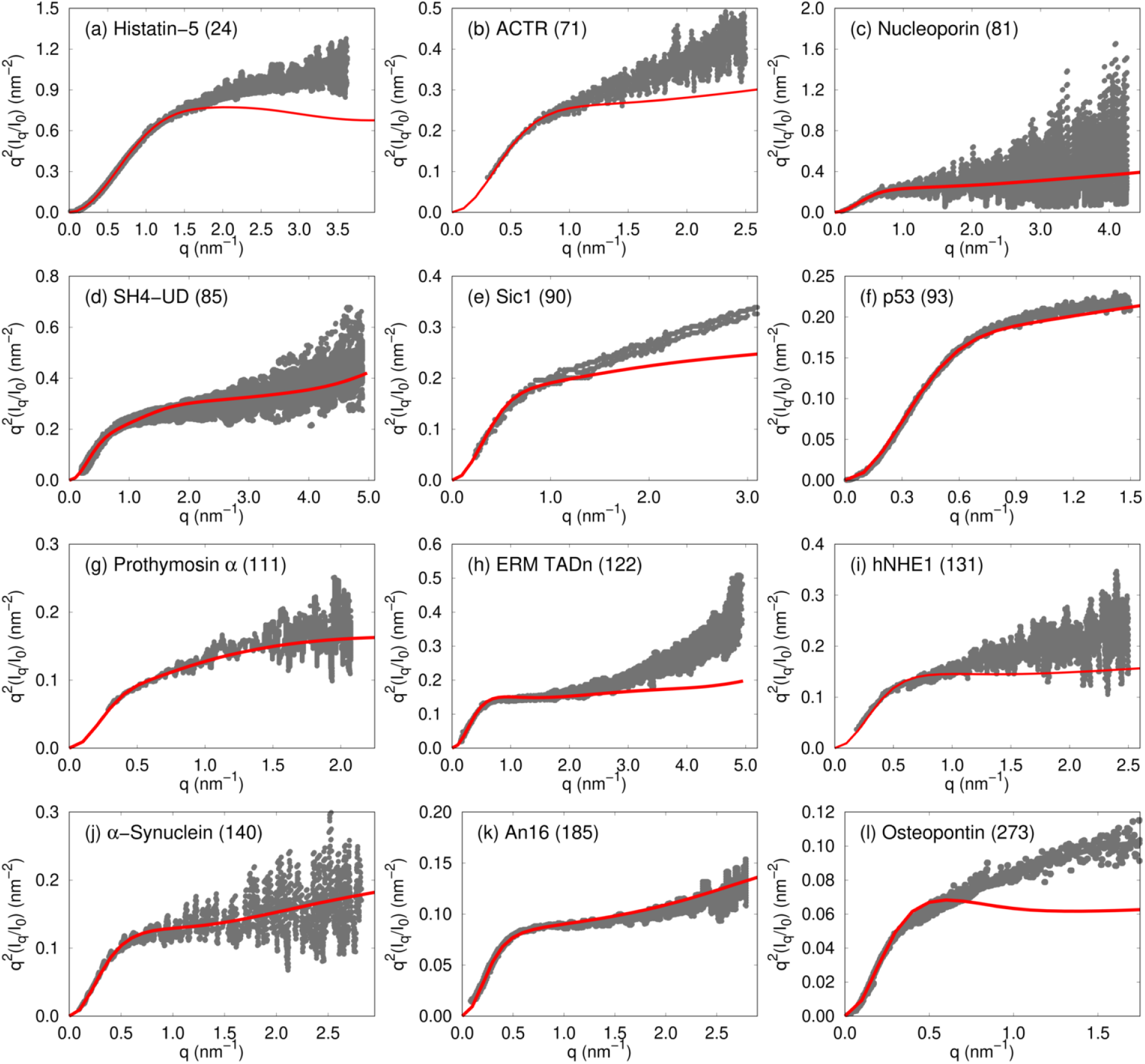
The Kratky plots for twelve IDP sequences. The values of *N*_*T*_ are in the parentheses. The gray points denote the experimental data, and the red curves report the simulated profiles. In almost all cases, the agreement between the simulations and experiment is excellent in the small *q* region.

**Figure S2:**
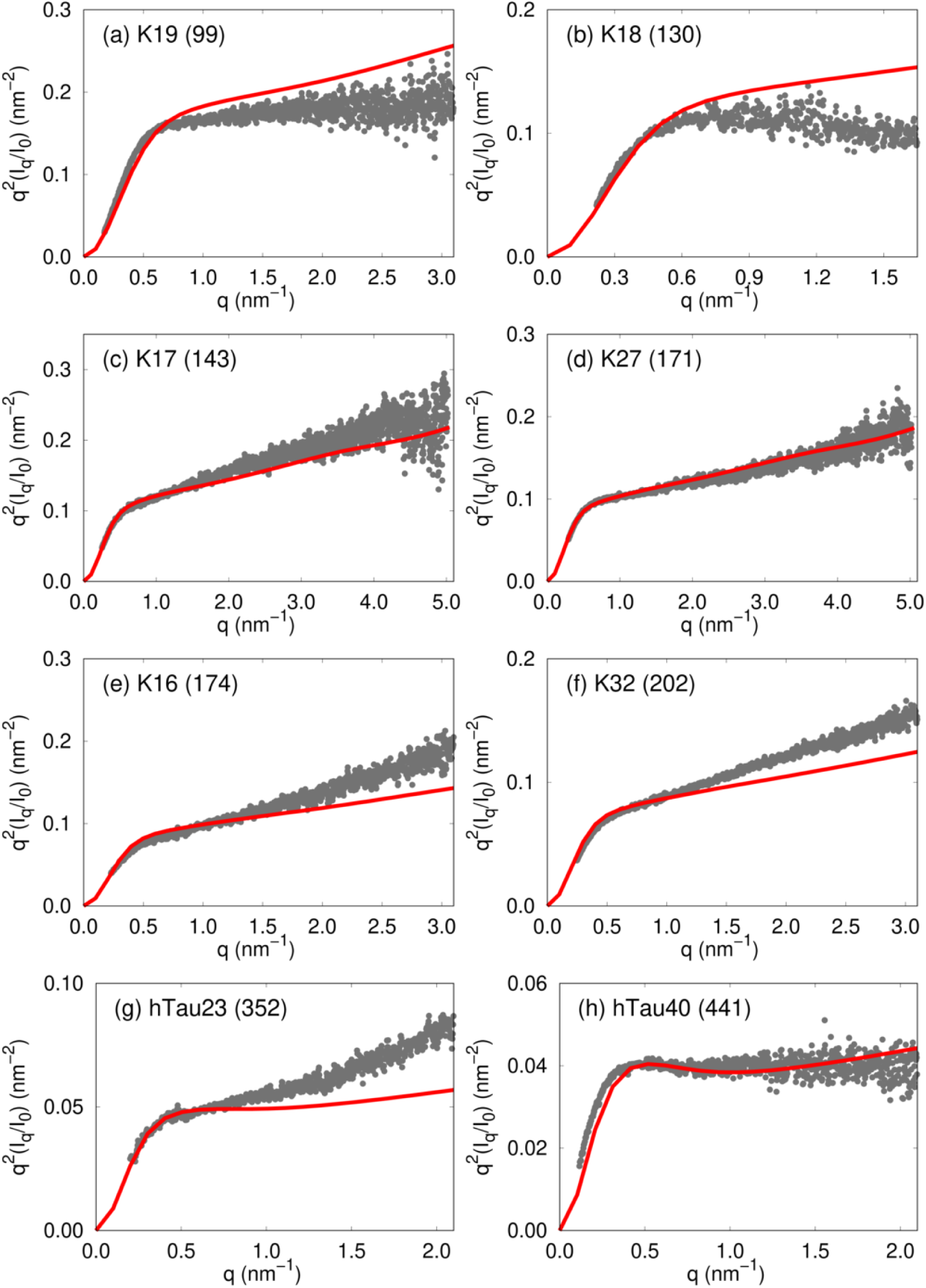
The Kratky plots for the Tau IDP sequences. The gray points denote the experimental data, and the red curves denote the simulated profiles. As in Fig. S1, the values of *N*_*T*_ are given in parentheses.

**Table S2:** Calculated *δ*^2^ between experimental and simulated SAXS profiles. The numbers in parentheses (*M*) represent the number of data points compared for the respective estimates.

| IDP | $(\delta^2\vert _{(qR_g)_{max}=3.0 \times 10^{-4}}) (M)$ | $(\delta^2\vert _{(qR_g)_{max}=5.0 \times 10^{-4}}) (M)$ |
| --- | --- | --- |
| Histatin-5 | 6.81 (22) | 79.42 (36) |
| ACTR | 2.08 (9) | 10.22 (17) |
| Nucleoporin | 1.81 (11) | 3.12 (19) |
| SH4-UD | 1.87 (9) | 0.91 (17) |
| Sic1 | 1.73 (8) | 2.54 (15) |
| p53 | 0.18 (10) | 0.41 (14) |
| Prothymosin $\alpha$ | 0.11 (5) | 0.70 (10) |
| ERM TADn | 2.05 (8) | 3.51 (15) |
| hNHE1 | 1.26 (7) | 2.18 (13) |
| α-Synuclein | 0.65 (7) | 0.81 (13) |
| An16 | 0.19 (5) | 0.34 (9) |
| Osteopontin | 0.53 (10) | 3.72 (14) |
| K19 | 1.02 (9) | 3.37 (15) |
| K18 | 1.13 (6) | 5.97 (12) |
| K17 | 0.01 (5) | 0.08 (11) |
| K27 | 0.18 (5) | 0.16 (10) |
| K16 | 0.05 (5) | 0.10 (10) |
| K32 | 0.03 (4) | 0.07 (9) |
| hTau23 | 0.02 (4) | 0.03 (7) |
| hTau40 | 0.08 (3) | 0.05 (6) |

In Eq. 4, *N* denotes the number of beads, *r*_*i*_ are the coordinates of bead *i, r*_*CM*_ is the centre-of-mass coordinate, and ⟨…⟩ denotes the ensemble average.

The correlation plot shown in Fig. S3 quantifies the high level of agreement between the experimental and simulated *R*_*g*_ values. The corresponding % errors in our prediction are quoted in Table S3. Using an expression similar to Eq. 3, we find that the relative error (*δ*^2^) between the experimental and simulated *R*_*g*_ values is 0.13.

**Figure S3:**
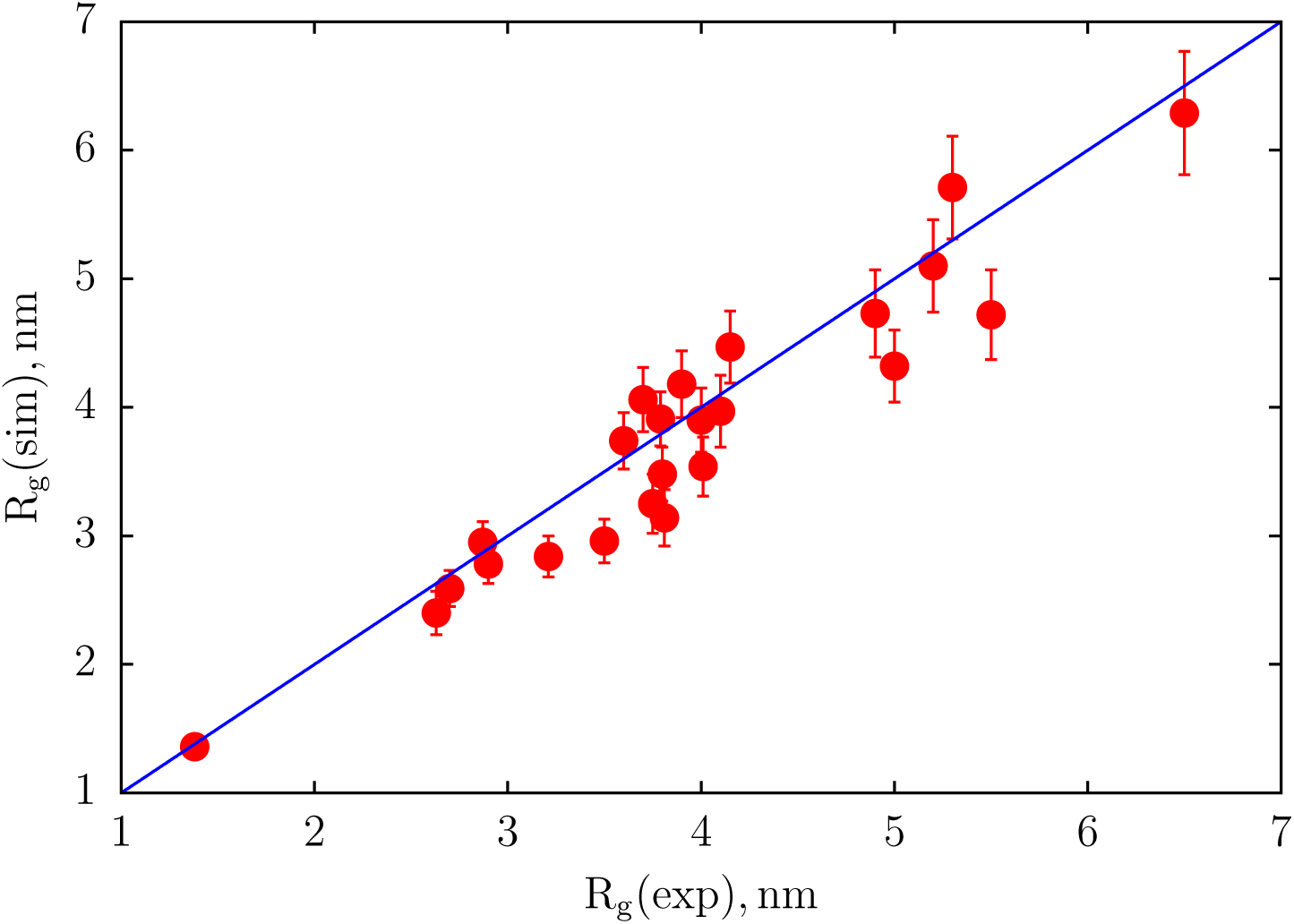
Comparison between the experimental *R*_*g*_ values and those obtained from simulations for the 24 IDP sequences. The blue solid line, which provides a guide to the eye, shows excellent agreement, especially when considering errors in experiments.

**Table S3:** Conformational properties of IDPs. The values of *R*_*g*_, *R*_*ee*_ and *R*_*h*_ are in nm. The numbers in parenthesis represent standard errors. Column 5 denotes the % error 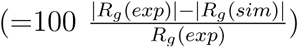 in our prediction of *R*_*g*_ values for the different IDPs. The reasons for large errors in the mean Δ and *S* are explained in the SI text.^*a*^ denotes the experimental value of *R*_*h*_ for the Sic1 sequence at 150 mM reported in Liu *et al.*^11^ ^*b*^ denotes the experimental *R*_*h*_ value for Prothymosin-*α* reported by Uversky *et al.*^12^ ^*c*^ denotes the experimental *R*_*h*_ value for *α*-Synuclein estimated by Paleologou *et al.*^13^

| IDP | $N_T$ | $R_g$ (exp) | $R_g$ (sim) | % error | $R_{ee}$ (sim) | $R_h$ (sim) | $\Delta$ | $S$ |
| --- | --- | --- | --- | --- | --- | --- | --- | --- |
| Histatin-5 | 24 | 1.38 | 1.36 | 1.5 | 3.09 | 1.32 | 0.40 (0.16) | 0.50 (0.34) |
| ACTR | 71 | 2.63 | 2.40 | 8.7 | 5.62 | 2.07 | 0.41 (0.19) | 0.49 (0.39) |
| Nucleoporin | 81 | 2.7 | 2.59 | 4.0 | 6.18 | 2.19 | 0.40 (0.19) | 0.49 (0.41) |
| SH4-UD | 85 | 2.9 | 2.78 | 4.1 | 6.55 | 2.30 | 0.43 (0.19) | 0.53 (0.43) |
| Sic1 | 90 | 3.21 | 2.84 | 11.5 | 6.77 | 2.36 (2.2)<sup>a</sup> | 0.42 (0.19) | 0.52 (0.40) |
| p53 | 93 | 2.87 | 2.95 | 2.8 | 7.08 | 2.43 | 0.42 (0.19) | 0.54 (0.40) |
| Prothymosin $\alpha$ | 111 | 3.79 | 3.91 | 3.2 | 9.37 | 2.92 (3.1)<sup>b</sup> | 0.51 (0.19) | 0.72 (0.45) |
| ERM TADn | 122 | 3.81 | 3.14 | 17.5 | 7.37 | 2.63 | 0.38 (0.19) | 0.45 (0.39) |
| hNHE1 | 131 | 3.75 | 3.25 | 13.3 | 7.45 | 2.69 | 0.39 (0.19) | 0.48 (0.39) |
| α-Synuclein | 140 | 4.00 | 3.54 | 11.7 | 8.16 | 2.89 (2.82)<sup>c</sup> | 0.40 (0.19) | 0.47 (0.40) |
| An16 | 185 | 5.0 | 4.32 | 13.5 | 10.24 | 3.38 | 0.43 (0.19) | 0.54 (0.41) |
| Osteopontin | 273 | 5.5 | 4.72 | 14.2 | 10.08 | 3.82 | 0.40 (0.19) | 0.52 (0.40) |
| K19 | 99 | 3.5 | 2.96 | 15.4 | 7.00 | 2.44 | 0.43 (0.19) | 0.54 (0.41) |
| K18 | 130 | 3.8 | 3.48 | 8.4 | 8.24 | 2.80 | 0.43 (0.19) | 0.54 (0.42) |
| K17 | 143 | 3.6 | 3.74 | 3.8 | 8.89 | 2.98 | 0.43 (0.19) | 0.54 (0.42) |
| K10 | 167 | 4.0 | 3.90 | 2.5 | 9.08 | 3.13 | 0.41 (0.19) | 0.50 (0.41) |
| K27 | 171 | 3.7 | 4.06 | 9.7 | 9.64 | 3.20 | 0.43 (0.19) | 0.55 (0.42) |
| K16 | 174 | 3.9 | 4.18 | 7.2 | 9.89 | 3.28 | 0.42 (0.19) | 0.51 (0.40) |
| K25 | 185 | 4.1 | 3.97 | 3.2 | 9.03 | 3.22 | 0.36 (0.18) | 0.41 (0.37) |
| K32 | 202 | 4.15 | 4.47 | 7.7 | 10.57 | 3.49 | 0.42 (0.19) | 0.52 (0.42) |
| K23 | 254 | 4.9 | 4.73 | 3.5 | 10.75 | 3.75 | 0.38 (0.19) | 0.44 (0.39) |
| K44 | 283 | 5.2 | 5.10 | 1.9 | 11.63 | 4.01 | 0.40 (0.19) | 0.49 (0.40) |
| hTau23 | 352 | 5.3 | 5.71 | 7.7 | 12.79 | 4.44 | 0.42 (0.19) | 0.53 (0.40) |
| hTau40 | 441 | 6.5 | 6.29 | 3.2 | 14.11 | 4.88 | 0.42 (0.19) | 0.54 (0.41) |

### Comparisons with results from all-atom force fields

In this section, we compare SOP-IDP simulation results for the SAXS profiles to available results obtained using simulations based on atomically detailed force fields. In the last few years, atomic detailed simulations, with vastly different force fields, have been used to calculate the SAXS profiles for the seventy one residue ACTR^8^ and the 24-residue RS peptide (FASTA sequence GAMGPSYGRSRSRSRSRSRSRSRS).^14^ For ACTR,^8^ the simulated *I*(*q*) and the radius of gyration (*R*_*g*_), which were calculated by adjusting the Lennard-Jones interaction strength between the oxygen atom of water and the heavy atoms on the protein within the AMBER force field, gave improved results relative the the Amber ff03 force field. As noted by the authors^8^ the overall dimension of ACTR is still more compact relative to the experimentally measured *R*_*g*_. However, simulations using the same force field as before^8^ performed in a larger simulation box have lead to *I*(*q*) (Best, personal communication) for ACTR that is in as excellent agreement with predictions based on SOP-IDP simulations for ACTR. (see Fig. 1b in the main text).

The experimental measurements of the SAXS profile for the RS peptide^15^ have been used to compare and validate the state-of-the-art all-atom force fields.^14,15^ Using an entirely different force field (TIP4P-D water model, adjustment of residue specific parameters, and addition of corrections to H bond interactions) Wu, Jiang, and Wu (WJW) arrived at the RSFF2+/TIP4P-D potential, here referred to as the WJW model. The WJW force field was used to calculate the SAXS profiles for the 24 residue Histatin-5 (see Fig. 1a in the main text) and the RS peptide. Comparison of these results with experiments showed excellent agreement with experimental *I*(*q*) (see Fig. S7 in Wu *et al.*^14^). Note that the force fields used in these studies^8,14–16^ are very different. For example, both in^14,16^ the H-bond between the backbone carbonyl oxygen and amide hydrogen is unusually short (0.15 nm versus the usual ∼ 0.2 nm in AMBER force fields) and the corresponding Lennard-Jones interaction is considerably stronger (∼ 0.3 kcal/mole versus 0.057 kcal/mole in AMBER). There are many other parameters that also vary greatly between the different force fields.

**Figure S4:**
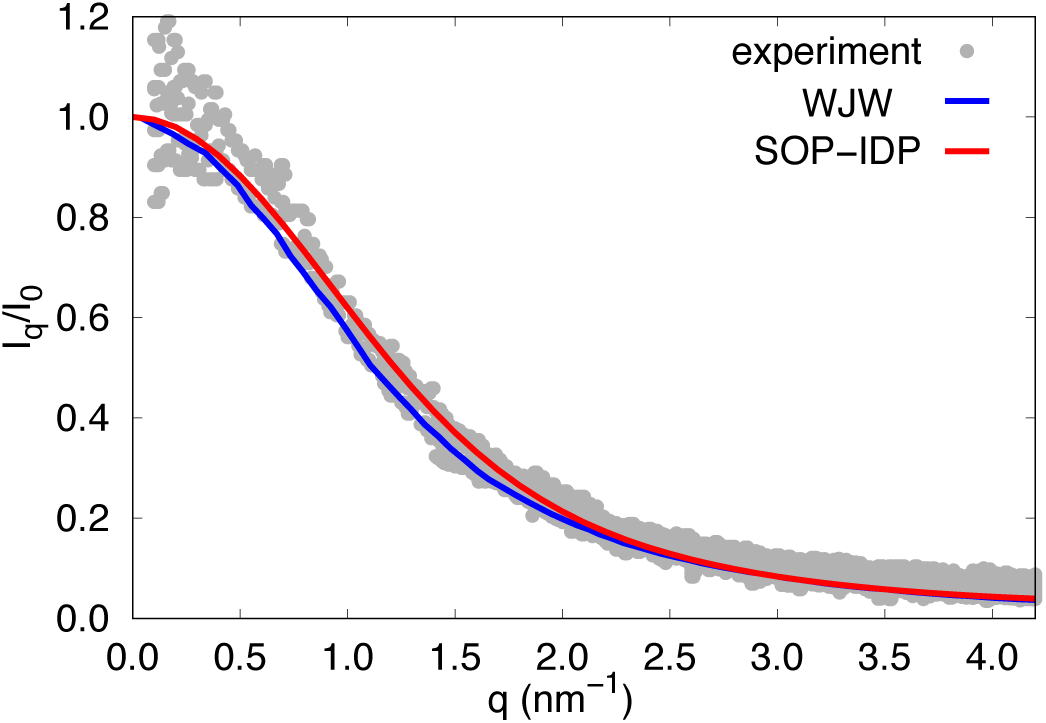
Top: Comparison of simulated SAXS profiles for the RS peptide using the SOP-IDP model with experiment,^15^ and MD simulations by Wu *et. al.* (RSFF2+).^14^

In the following, we briefly compare results from simulations using the SOP-IDP model for the RS peptide, with results from all-atom simulation models, and experiment. The results of the SAXS experiments for the RS peptide were reported by Grubmüller *et. al.* at 298 K temperature, neutral pH, 100 mM NaCl, and 50 mM Na-phosphate buffer. The experimental value of *R*_*g*_ is 1.262*±*0.007 nm.^15^ They also benchmarked the performances of multiple state-of-the-art peptide force-fields including AMBER, OPLS, and CHARMM variants (for details please refer to Table 1 in Grubmüller *et al.*^15^) against the experimental observations (Recent comparison of the performances of different force fields may be found in^14,16^). The large-scale simulations, using replica-exchange molecular dynamics at temperatures between 298 K and 450 K, were performed at 150 mM NaCl concentration.^15^ The best agreement with the experimental *R*_*g*_ was obtained with the CHARMM 22* force-field (1.265*±*0.007 nm) in conjunction with CHARMM-modified TIP3P water. The same peptide force-field when used in conjunction with the dispersion-corrected TIP4P-D water model, ^14,17^ however resulted in a somewhat larger *R*_*g*_ of ∼1.4 nm. Several other combinations of peptide and water force fields yielded a wide range of *R*_*g*_ values ∼(1.0–1.5) nm.^15^ The RS peptide was also used to validate the WJW force-field, ^14^ which was optimized based on the AMBER-ff99SB but with a different water model and adjustments to many other parameters. The *R*_*g*_ value was obtained to be (1.32*±*0.005 nm), also using replica-exchange molecular dynamics simulations, and at salt concentration adequate for neutralization of the peptide charges.

Using the SOP-IDP model at 150 mM salt concentration, we obtained the *R*_*g*_ value of 1.293*±*0.07 nm) for the RS peptide, which is in excellent agreement (a mere 2.5 % difference) with the experimental measurement. In Fig. S4 the SAXS profiles obtained from our simulations of the RS peptide using the SOP-IDP model is compared with the experimental SAXS profile,^15^ and simulated *I*(*q*) using the WJW force-field.^14^ The latter data are extracted from the corresponding references using the online tool *WebPlotDigitizer*.^18^ As can clearly be seen, simulated profiles from both the models are within the dispersion of experimental data. This comparison also illustrates that one could construct several force-fields (both atomically detailed as well as coarse-grained models), which could yield comparable results for small IDPs. The challenge is to *predict* results for IDPs of arbitrary length and sequences. Comparison with experiments will ultimately be the sole test of accuracy.

**Figure S5:**
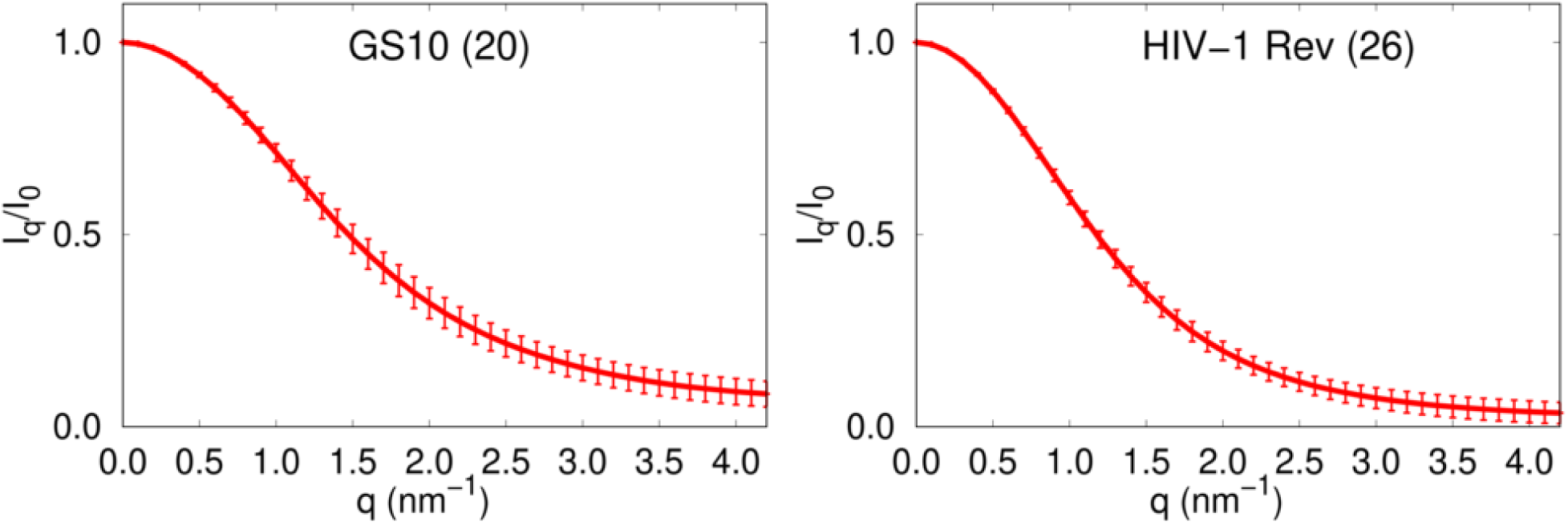
Predicted SAXS profiles for the GS10 peptide and the argininine rich motif of HIV-1 Rev (see text for sequences and simulation conditions) using the SOP-IDP model. The numbers in parentheses represent sequence lengths.

### Predictions using the SOP-IDP model

We also extended the SOP-IDP simulations to two additional disordered sequences, the charge-neutral GS10 (FASTA sequence GSGSGSGSGSGSGSGSGSGS) and the arginine rich motif of HIV-1 Rev (FASTA sequence GAMATRQARRNRRRRWRERQRAAAAR). To the best of our knowledge, experimental SAXS profiles are not available for these peptides. For these sequences, WJW reported reported the *R*_*g*_ values ∼(.936*±*0.05 nm) for GS10 and ∼(1.49*±*0.04 nm) for HIV-1 Rev^14^ (data extracted digitally from pink colored bar graph in Figure S6 in Wu *et al.*^14^). The corresponding values from simulations using the SOP-IDP model at 298 K simulation temperature are (1.027*±*0.08 nm) and (1.465*±*0.096 nm) for GS10 and HIV-1, respectively. In these two cases the *R*_*g*_ values predicted using simulations with the state of the art WJW force field and the SOP-IDP simulations are in excellent agreement. The simulations for the HIV-1 Rev peptide using the SOP-IDP model were performed at effective salt concentration of 10 mM (simulations by Wu *et al.* were reported at salt concentration adequate for neutralization of the peptide charges). In Fig. S5 we plot the simulated SAXS profiles. For both these peptides the calculated *I*(*q*) profiles serve as testable predictions.

A note concerning achieving good agreement between simulations and experiments is in order. For the three small IDPs (20 residue GS10, 24 residue RS peptide, and the 26 residue HIV-1 Rev) the *R*_*g*_ values can be readily calculated using the fit *R*_*g*_ = 0.2*N* ^0.588^ given in the caption to Fig. 3a in the main text. This Flory formula gives *R*_*g*_ values 1.164 nm, 1.296 nm, and 1.358 nm, for GS10, RS peptide, and HIV-1 Rev respectively. These theoretical results do not differ significantly from the simulation results quoted above, which shows that in order to assess the accuracy of the force fields, at any level of coarse-graining, for IDPs or globular proteins one has to compare predictions with directly measurable quantities, such as *I*(*q*) as a function of *q* and heat capacity.

### Distributions of shape parameters

In Figures S6–S9, we show the distributions of shape parameters Δ and *S* (see Eq. 5 in main text) for all 24 IDP sequences. Remarkably, the results in Figures S6–S9 show that *P* (Δ) and *P* (*S*) are broad, with large dispersions. This is due to the plasticity of conformations sampled by the IDPs, which is further corroborated using hierarchical clustering of the IDP conformations (as discussed in the main text.) The large dispersions result in substantial standard errors (Table S3), which merely shows that the average Δ and *S* do not elucidate the granularity of the IDP conformations.

**Figure S6:**
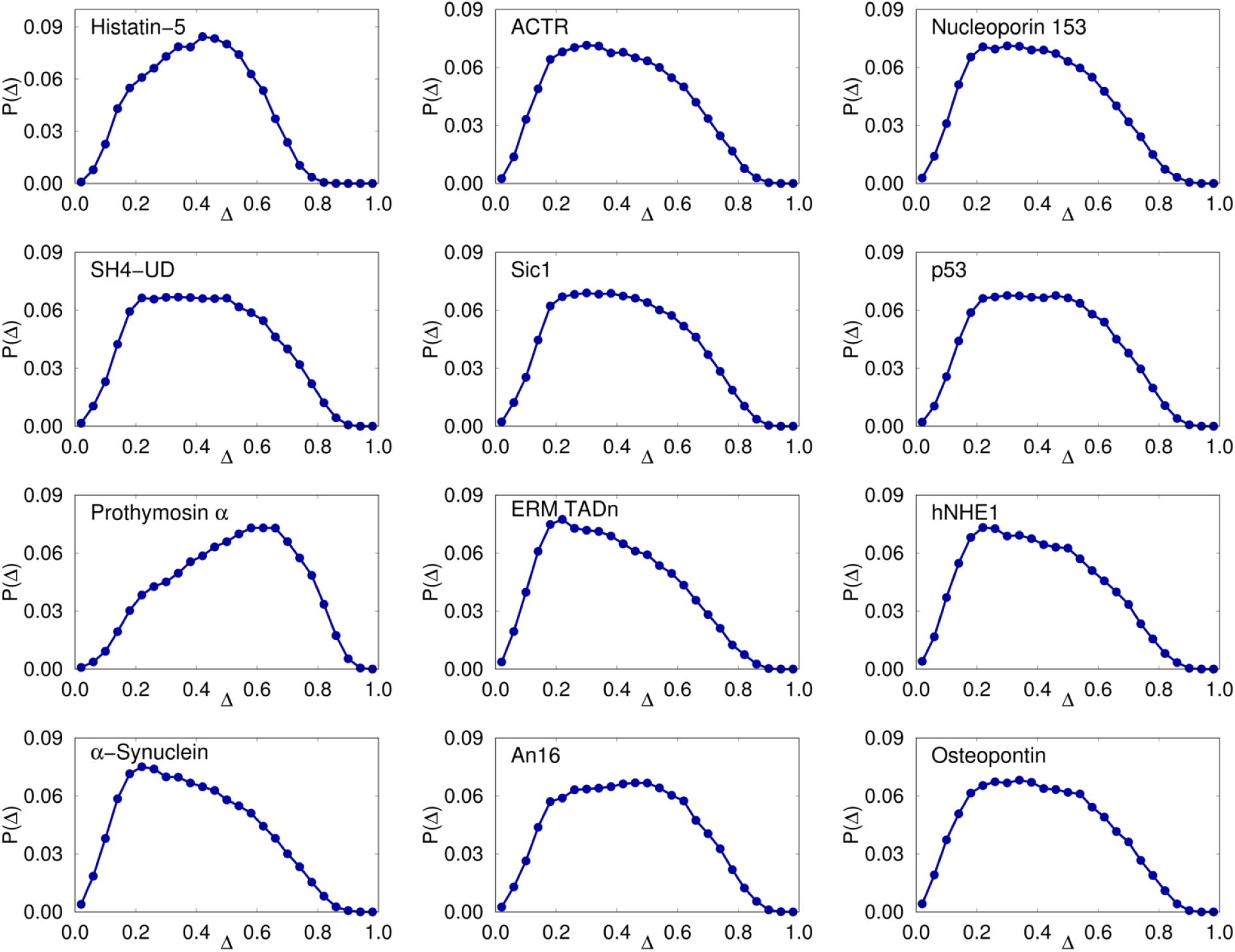
Distributions of the shape parameter Δ (Eq. 6 in the main text), for all non-Tau IDP sequences listed in Table S3.

**Figure S7:**
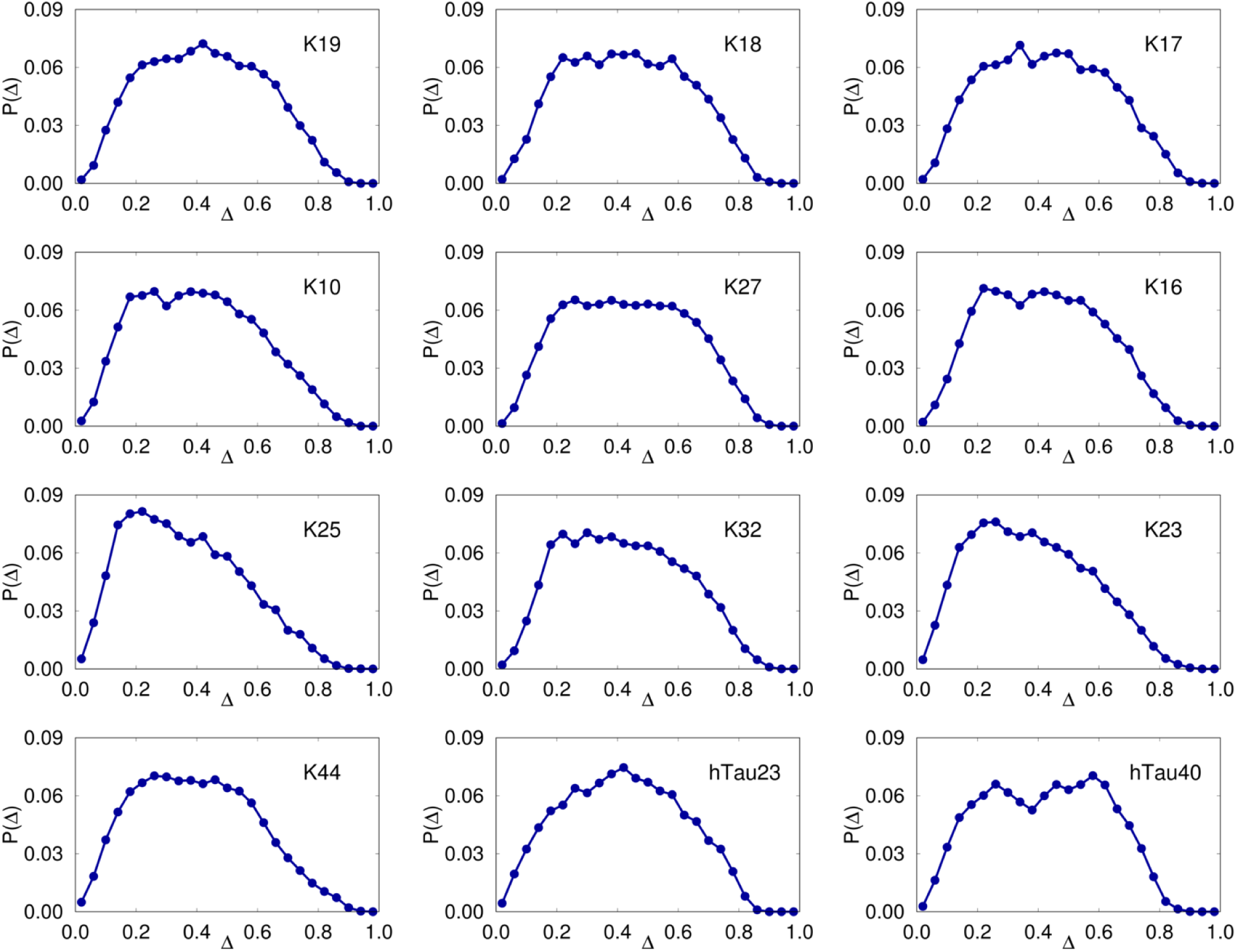
Same as Figure S6, except the results are for the Tau protein constructs.

**Figure S8:**
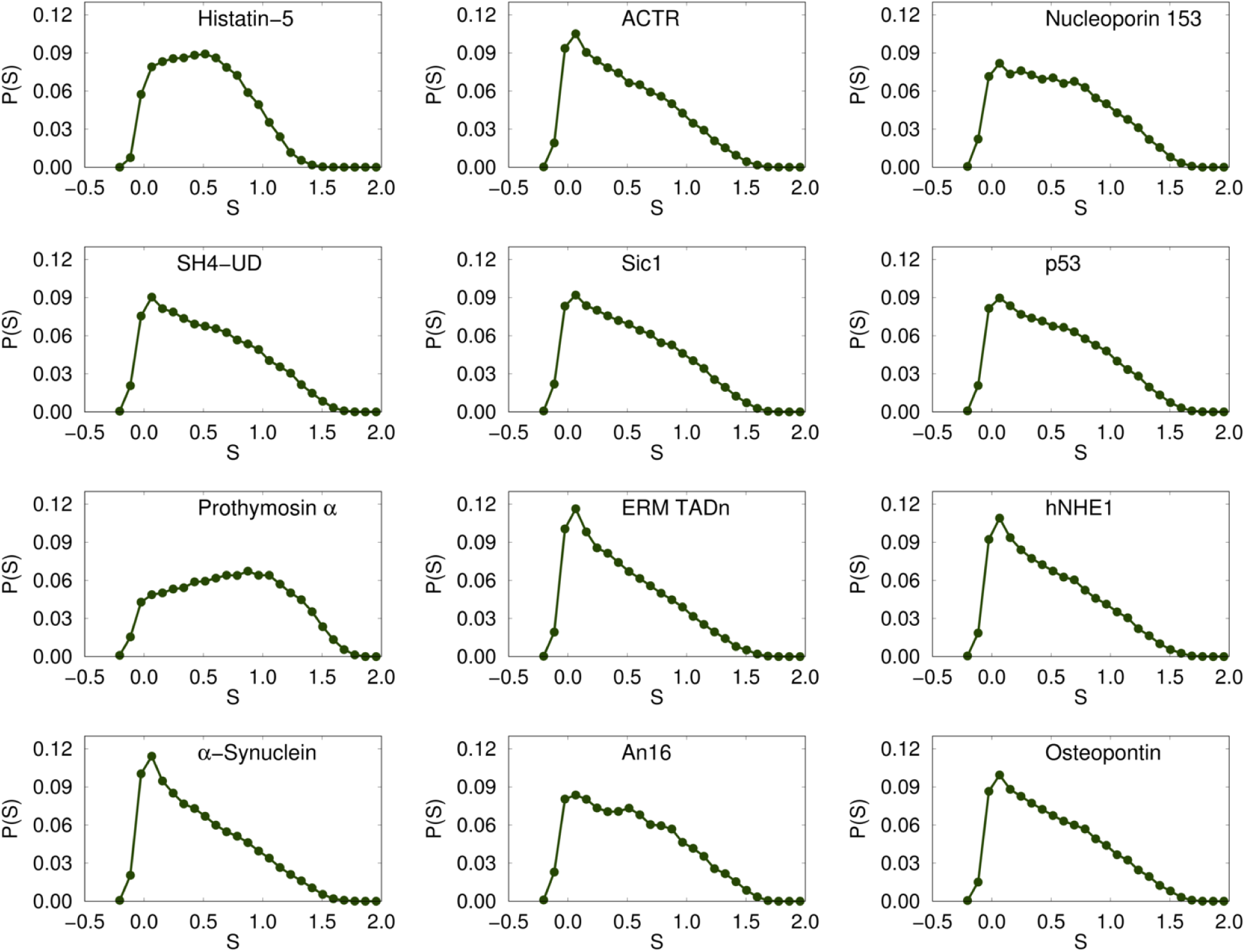
Distributions of the shape parameter *S* (Eq. 6 in the main text), for the twelve non-Tau IDP sequences.

**Figure S9:**
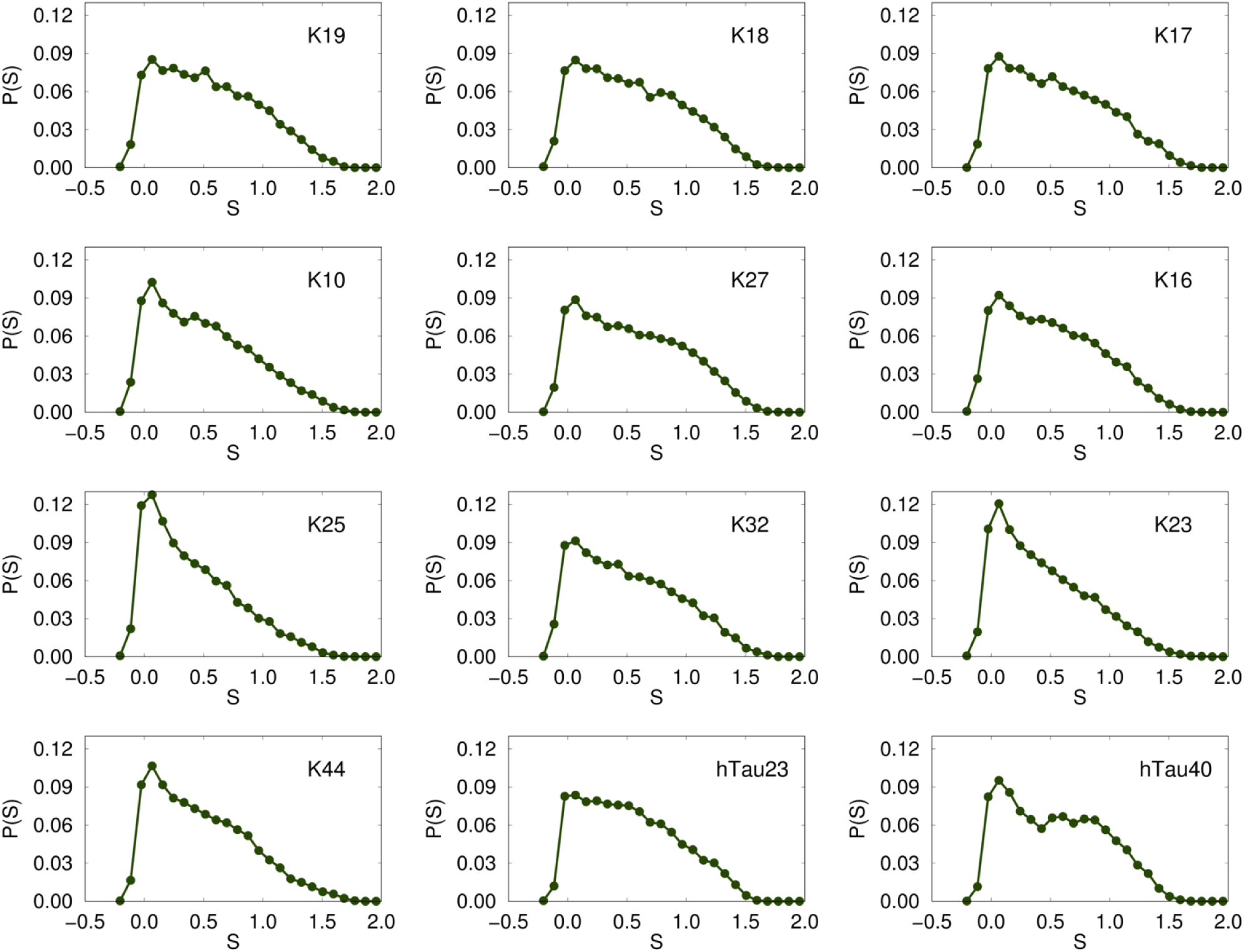
Same as Figure S8, except the results are for the Tau protein constructs.

### Distribution of Radius of Gyration, *R*_*g*_

In Figures S10 and S11, we show the *R*_*g*_ distributions for all the IDP sequences, which can be computed only in simulations. From experiments, one can only obtain the distance distribution functions by inverse Fourier transform of *I*(*q*)s. Nevertheless, these results are useful in assessing the extent of conformational fluctuations, and for benchmarking our simulations against other force fields.

**Figure S10:**
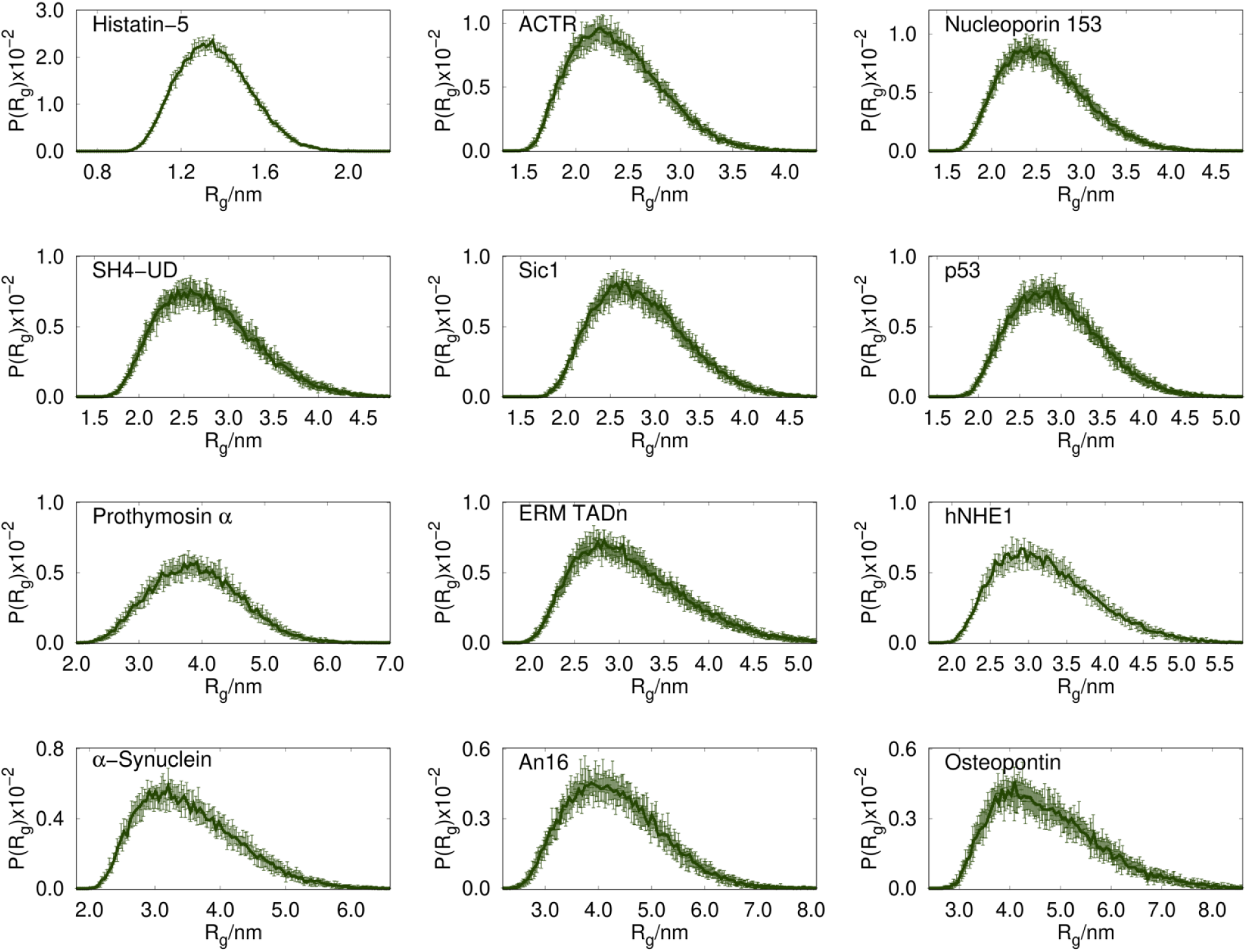
Distribution of *R*_*g*_ for all the non-Tau IDP sequences listed in Table S3.

**Figure S11:**
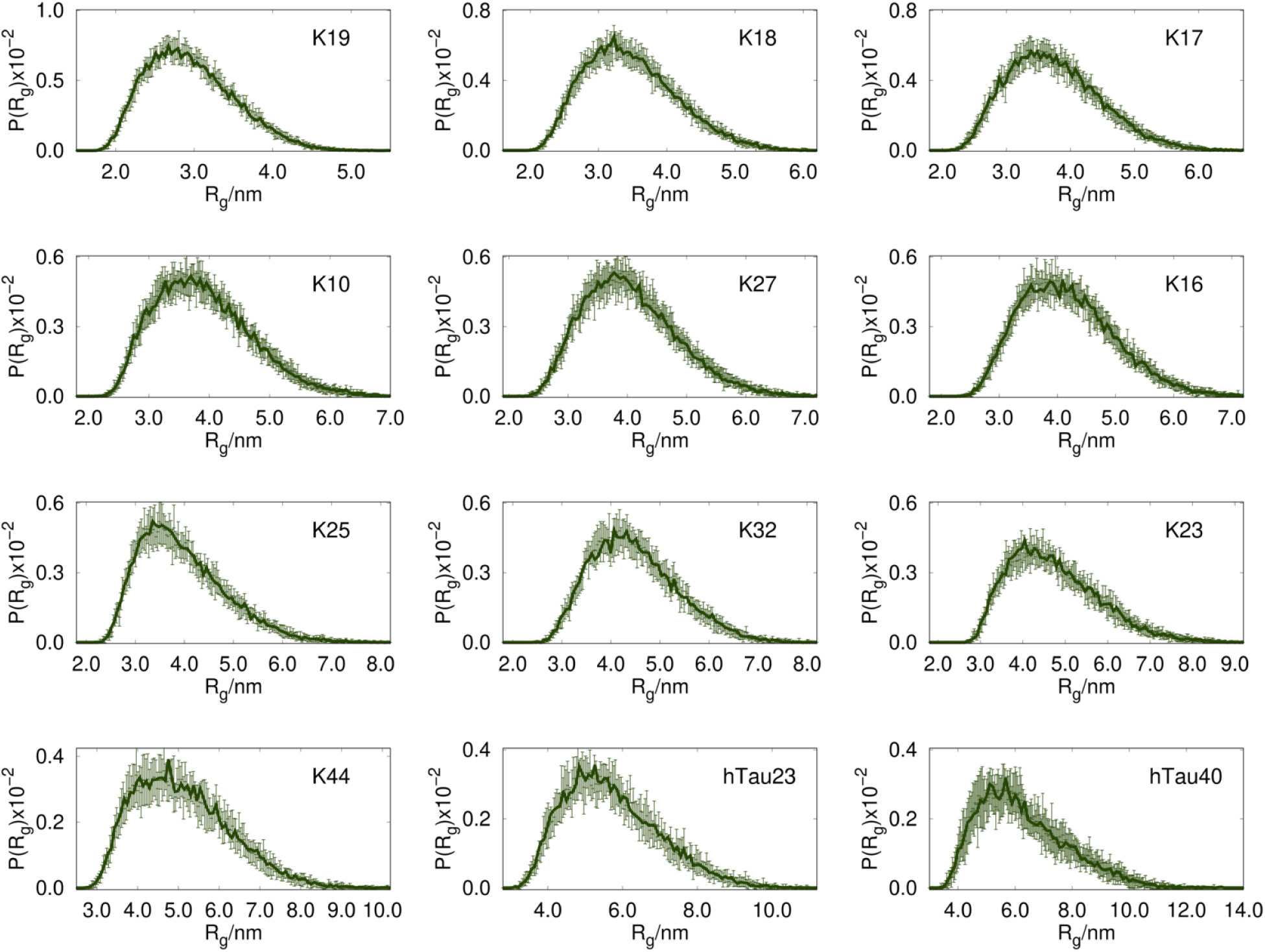
Distribution of *R*_*g*_ for all the Tau IDP sequences listed in Table S3.

### End-to-end (*R*_*ee*_) distance distributions, *P* (*R*_*ee*_)s

In Figures S12 and S13, we show the *R*_*ee*_ distributions for all 24 IDP sequences, and compare them with rigorous theoretical curves for the random coil-like behavior, and the Gaussian chain. The theoretical results for the random coil-like and Gaussian chain is given by^19^ 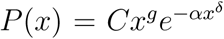 for *N_T_* >> 1, where 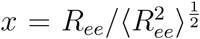 and *C* is a normalization constant. The exponent *δ* = 1*/* (1 − *v*) (with *v* ≈ 0.588 for a random coil and *ν* = 0.5 for a Gaussian chain) accounts for the decay of *P* (*x*) for *x* > 1. The correlation hole exponent *g* is ≈ 0.28 for a random coil and represents the reduced probability of finding the end of the chains in contact in the presence of repulsive interactions. For a Gaussian chain, *g* = 0. The conditions 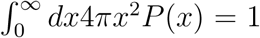 and 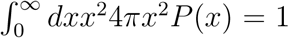 allow the determination of *C* and *α*.^19^ For a Gaussian chain *C* = (3*/*2*π*)^3^*/*^2^ and *α* = 1.5, whereas *C ≈* 0.278 and *α ≈* 1.206 are approximate results for a random coil. In addition, when *N*_*T*_ >> 1, the effects of side chains on *P* (*R*_*ee*_) is unimportant, whereas for finite *N*_*T*_, as is the case for IDPs considered here, they can be significant.

Interestingly, for the Tau protein constructs, the *P* (*R*_*ee*_) distributions in the range 99 ≤ *N*_*T*_ ≤ 174 (first six panels in Figure S13) are well described by the *P* (*R*_*ee*_) corresponding to the theoretical random coil-like behavior, while in the range 185 ≤ *N*_*T*_ ≤ 441 the distributions coincide with the predictions based on the Gaussian chain (last four panels in Figure S13). This is an example where the apparent solvent quality changes upon increasing *N*_*T*_.

**Figure S12:**
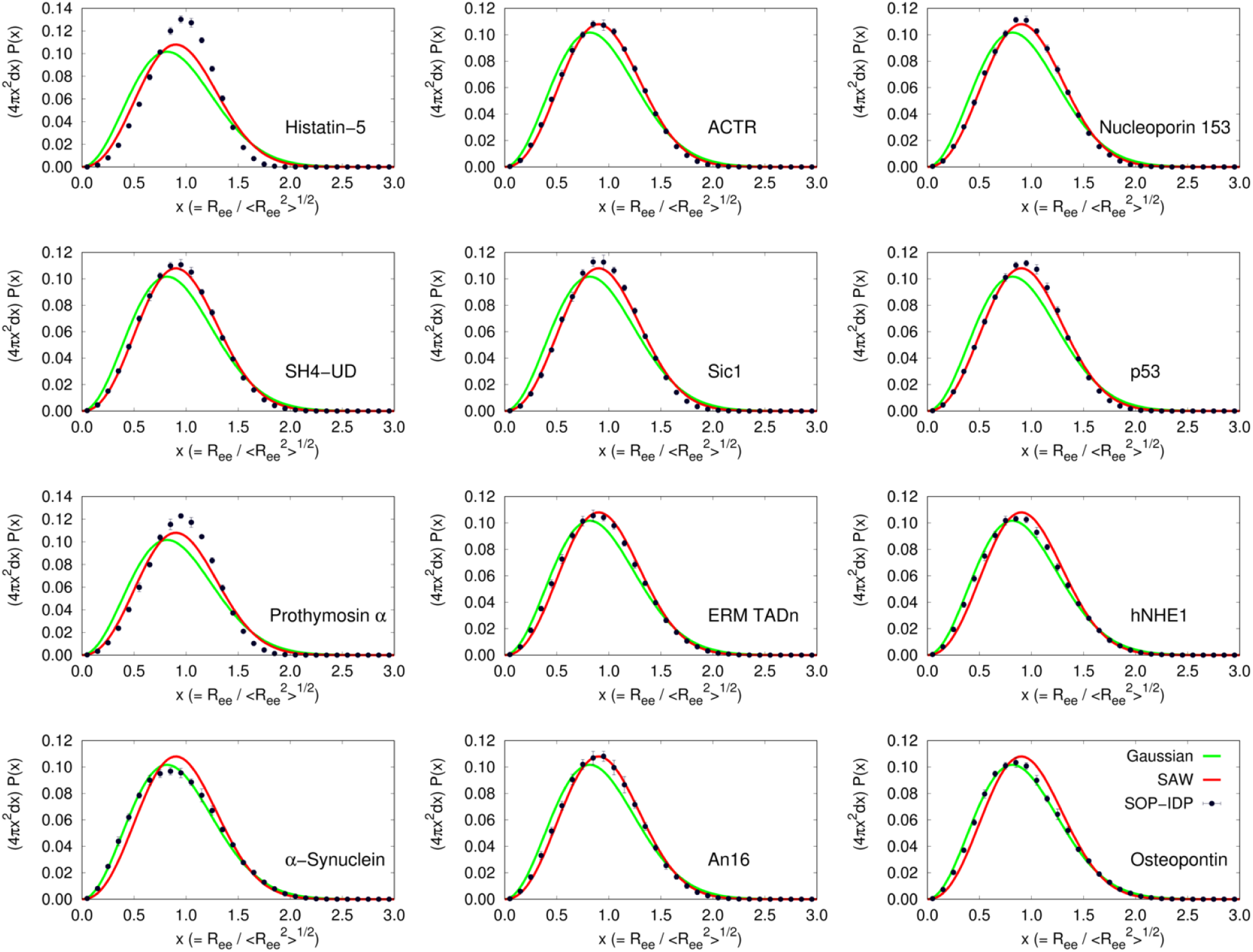
(black points) Plots for *R_ee_* (scaled by 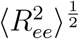 distance distributions for non-Tau IDP sequences. Theoretical distributions for random coil-like behavior and Gaussian chains are shown in red and green, respectively.

**Figure S13:**
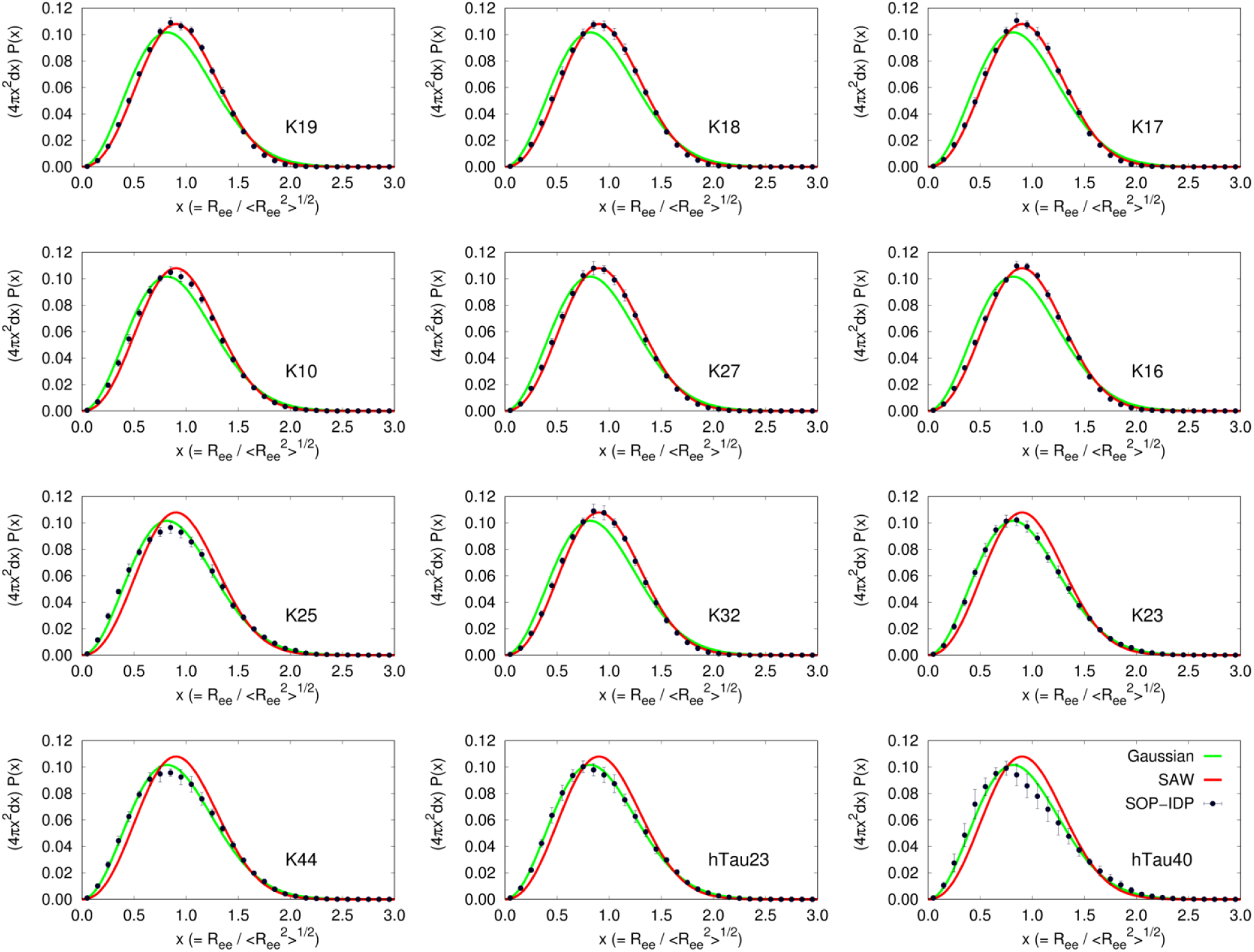
Same as Figure S12, except the results are for for Tau protein constructs. Theoretical distributions for random coil-like behavior and Gaussian chains are shown in red and green, respectively.

### Sequence compositional properties

Inspired by sequence variables that naturally arise in theory of polyampholytes (PAs), ^20,21^ the biophysical properties of IDPs are often characterized on the basis of sequence compositional properties such as the fraction of positively / negatively charged residues (*f*_+_ */ f*_−_), net fraction of charged residues (*f*_+_ + *f*_−_), as well as quantities related to charge asymmetry, 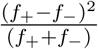.^22^ Based on these variables it is natural to construct the plausible phases that a given IDP might adopt under ambient conditions. It is known from polyelectrolyte ^23^ and PA^20^ theories that the phases could be readily altered by external conditions such as changing salt concentration. Clearly, the external conditions are not encoded in *f*_+_ and *f*_−_. Nevertheless, the PA type variable are useful in anticipating the states of IDP. The PA-like properties for the IDP sequences are tabulated in Table S4 for all the simulated sequences. Although such quantities may be relevant in qualitative descriptions, their use in inferring conformational properties of specific IDP sequences is very limited. There is no obvious distinction in the compositional properties among the diverse set of IDPs studied in this work. While they span a wide range in their respective compositional properties, an understanding of their heterogeneous conformational ensembles necessarily requires explicit simulations, as discussed in detail in the main text.

**Table S4:** Sequence compositional properties of the studied IDP sequences.

| IDP | $f_+$ | $f_-$ | $f_+ + f_-$ | $(f_+ - f_-)^2 / (f_+ + f_-)$ |
| --- | --- | --- | --- | --- |
| Histatin-5 | 0.29 | 0.08 | 0.38 | 0.116 |
| ACTR | 0.08 | 0.18 | 0.27 | 0.036 |
| Nucleoporin 153 | 0.00 | 0.00 | 0.00 | 0.00 |
| SH4-UD | 0.14 | 0.07 | 0.21 | 0.024 |
| Sic1 | 0.12 | 0.00 | 0.12 | 0.122 |
| p53 | 0.02 | 0.18 | 0.20 | 0.127 |
| Prothymosin $\alpha$ | 0.09 | 0.49 | 0.58 | 0.273 |
| ERM TADn | 0.13 | 0.18 | 0.31 | 0.008 |
| hNHE1 | 0.11 | 0.20 | 0.31 | 0.023 |
| α-Synuclein | 0.11 | 0.17 | 0.29 | 0.011 |
| An16 | 0.03 | 0.0 | 0.03 | 0.032 |
| Osteopontin | 0.15 | 0.25 | 0.41 | 0.024 |
| K19 | 0.20 | 0.09 | 0.29 | 0.042 |
| K18 | 0.20 | 0.08 | 0.28 | 0.047 |
| K17 | 0.20 | 0.07 | 0.27 | 0.064 |
| K10 | 0.18 | 0.10 | 0.28 | 0.021 |
| K27 | 0.21 | 0.08 | 0.29 | 0.060 |
| K16 | 0.20 | 0.07 | 0.27 | 0.064 |
| K25 | 0.17 | 0.13 | 0.30 | 0.005 |
| K32 | 0.21 | 0.08 | 0.28 | 0.061 |
| K23 | 0.16 | 0.13 | 0.29 | 0.004 |
| K44 | 0.18 | 0.12 | 0.30 | 0.014 |
| hTau23 | 0.17 | 0.12 | 0.29 | 0.011 |
| hTau40 | 0.16 | 0.13 | 0.29 | 0.011 |

### Contact maps reveal deviations from random coil-like behavior

As discussed in the main text, deviation from random-coil like behavior for certain IDPs, particularly the tau constructs, can be gleaned from their difference contact maps. In Figure S14, we show that the WT Tau sequence (hTau40) contains locally compact region, with enhanced contacts. This region is highlighted using a zoomed-in view of the contact map. The segment with locally enhanced contacts is also present in K25 (see Figure 8 in main manuscript for a zoomed-in view, and below for full range), and the K23, K44, and hTau23 constructs (see below).

**Figure S14:**
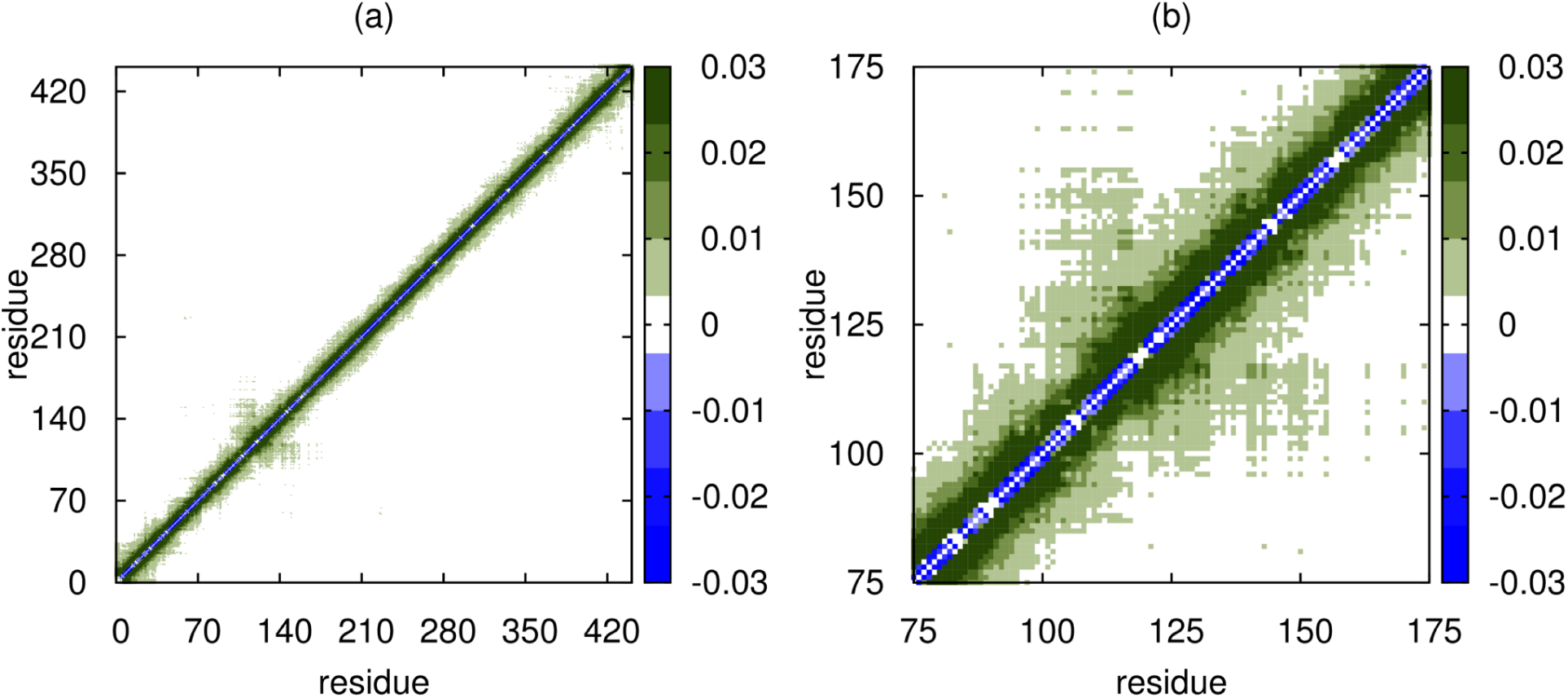
Difference contact map (defined in Eq.1 in the main text) between the one obtained using SOP-IDP simulations and the corresponding contact map for a Flory random coil for (a) the full length Tau protein (hTau40). (b) Zoomed-in view, highlighting the region with enhanced contacts.

**Figure S15:**
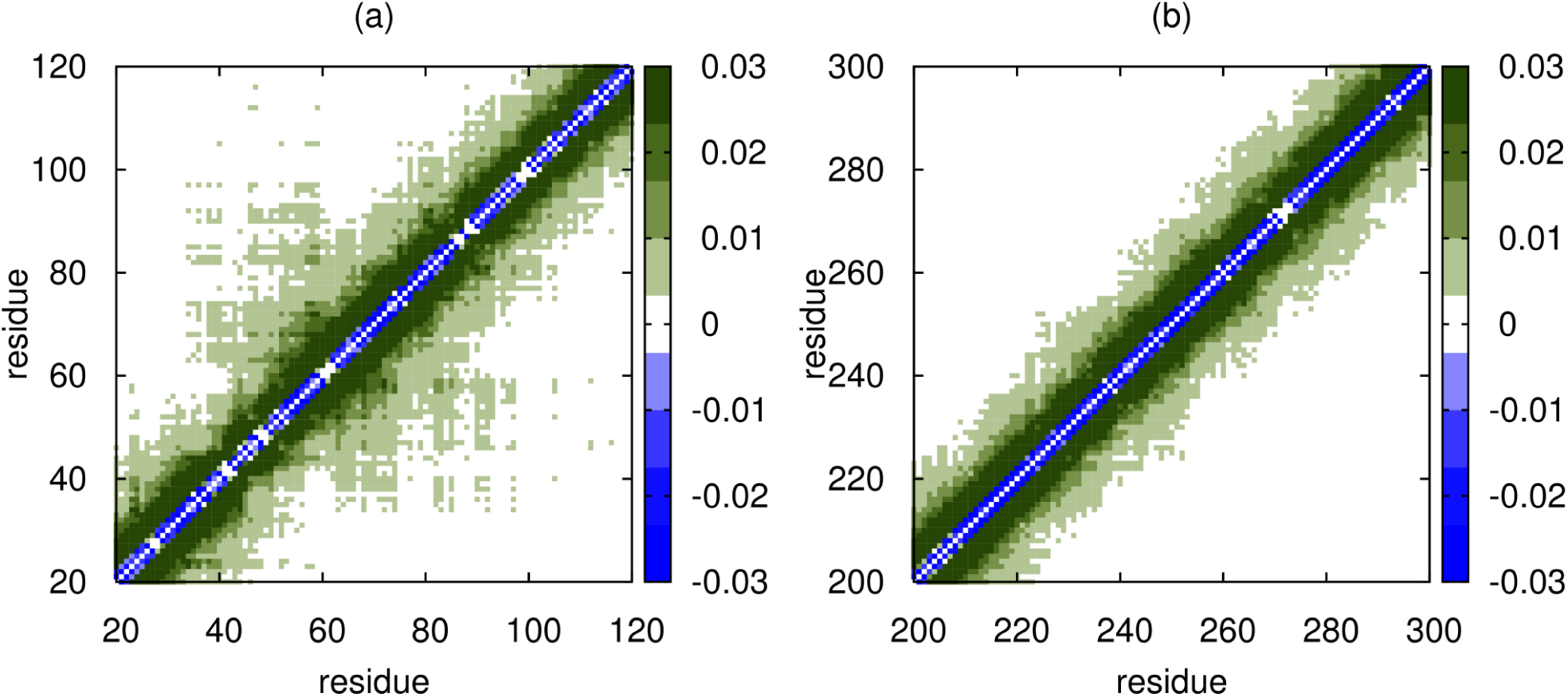
Same as Figure S14 except this if for the hTau23 sequence (a). Zoomed-in view for residues 200-300 of hTau40, shown for comparison (b).

**Figure S16:**
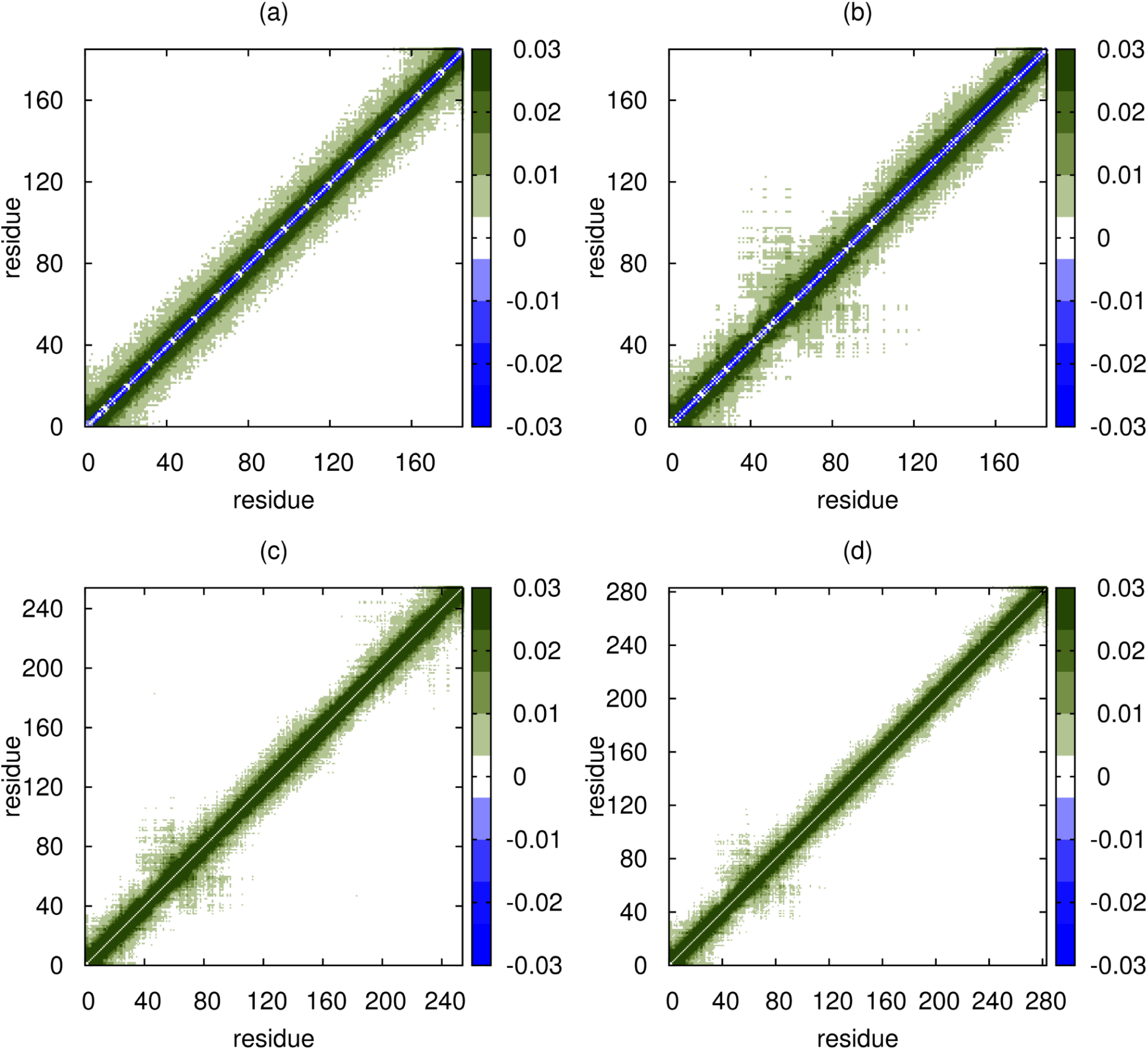
Difference contact maps (Eq. 1 in the main text) for (a) An16, (b) K25, (c) K23 and (d) K44, showing the full range in each case.

### Simulated sequences for the IDPs in the FASTA representation

We use one letter codes for the amino acids. In cases where the sequences were not explicitly provided in the experimental references, the sequences were obtained from database DisProt.^24^ As such, some sequences differ by ≤ 4 residues from the experimental sequence lengths. In such cases, the *R*_*g*_ values were scaled up/down following the appropriate 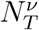 scaling.

- Histatin-5 DSHAKRHHGYKRKFHEKHHSHRGY
- ACTR GTQNRPLLRNSLDDLVGPPSNLEGQSDERALLDQLHTLLSNTDATGLEEIDRALGIPELVNQGQA LEPKQD
- Nucleoporin 153 GCPSASPAFGANQTPTFGQSQGASQPNPPGFGSISSSTALFPTGSQPAPPTFGTVSSSSQPPVFG QQPSQSAFGSGTTPNA
- SH4-UD MGSNKSKPKDASQRRRSLEPAENVHGAGGGAFPASQTPSKPASADGHRGPSAAFAPAAAEPKLFG GFNSSDTVTSPQRAGPLAGG
- Sic1 MTPSTPPRSRGTRYLAQPSGNTSSSALMQGQKTPQKPSQNLVPVTPSTTKSFKNAPLLAPPNSNM GMTSPFNGLTSPQRSPFPKSSVKRT
- p53 MEEPQSDPSVEPPLSQETFSDLWKLLPENNVLSPLPSQAMDDLMLSPDDIEQWFTEDPGPDEAPR MPEAAPPVAPAPAAPTPAAPAPAPSWPL
- Prothymosin *α* MSDAAVDTSSEITTKDLKEKKEVVEEAENGRDAPANGNAENEENGEQEADNEVDEEEEEGGEEEE EEEEGDGEEEDGDEDEEAESATGKRAAEDDEDDDVDTKKQKTDEDD
- ERM TADn MDGFYDQQVPFMVPGKSRSEECRGRPVIDRKRKFLDTDLAHDSEELFQDLSQLQEAWLAEAQVPD DEQFVPDFQSDNLVLHAPPPTKIKRELHSPSSELSSCSHEQALGANYGEKCLYNYCA
- hNHE1 MVPAHKLDSPTMSRARIGSDPLAYEPKEDLPVITIDPASPQSPESVDLVNEELKGKVLGLSRDPAK VAEEDEDDDGGIMMRSKETSSPGTDDVFTPAPSDSPSSQRIQRCLSDPGPHPEPGEGEPFFPKGQ
- *α*-Synuclein MDVFMKGLSKAKEGVVAAAEKTKQGVAEAAGKTKEGVLYVGSKTKEGVVHGVATVAEKTKEQVTN VGGAVVTGVTAVAQKTVEGAGSIAAATGFVKKDQLGKNEEGAPQEGILEDMPVDPDNEAYEMPSE EGYQDYEPEA
- An16 MHHHHHHPGAPAQTPSSQYGAPAQTPSSQYGAPAQTPSSQYGAPAQTPSSQYGAPAQTPSSQYGA PAQTPSSQYGAPAQTPSSQYGAPAQTPSSQYGAPAQTPSSQYGAPAQTPSSQYGAPAQTPSSQYG APAQTPSSQYGAPAQTPSSQYGAPAQTPSSQYGAPAQTPSSQYGAPAQTPSSQYV
- Osteopontin MRIAVICFCLLGITCAIPVKQADSGSSEEKQTLPSKSNESHDHMDDMDDEDDDDHVDSQDSIDSN DSDDVDDTDDSHQSDESHHSDESDELVTDFPTDLPATEVFTPVVPTVDTYDGRGDSVVYGLRSKS KKFRRPDIQYPDATDEDITSHMESEELNGAYKAIPVAQDLNAPSDWDSRGKDSYETSQLDDQSAE THSHKQSRLYKRKANDESNEHSDVIDSQELSKVSREFHSHEFHSHEDMLVVDPKSKEEDKHLKFR ISHELDSASSEVN
- K19 MQTAPVPMPDLKNVKSKIGSTENLKHQPGGGKVQIVYKPVDLSKVTSKCGSLGNIHHKPGGGQVE VKSEKLDFKDRVQSKIGSLDNITHVPGGGNKKIE
- K18 MQTAPVPMPDLKNVKSKIGSTENLKHQPGGGKVQIINKKLDLSNVQSKCGSKDNIKHVPGGGSVQ IVYKPVDLSKVTSKCGSLGNIHHKPGGGQVEVKSEKLDFKDRVQSKIGSLDNITHVPGGGNKKIE
- K17 MSSPGSPGTPGSRSRTPSLPTPPTREPKKVAVVRTPPKSPSSAKSRLQTAPVPMPDLKNVKSKIG STENLKHQPGGGKVQIVYKPVDLSKVTSKCGSLGNIHHKPGGGQVEVKSEKLDFKDRVQSKIGSL DNITHVPGGGNKKIE
- K10 MQTAPVPMPDLKNVKSKIGSTENLKHQPGGGKVQIVYKPVDLSKVTSKCGSLGNIHHKPGGGQVE VKSEKLDFKDRVQSKIGSLDNITHVPGGGNKKIETHKLTFRENAKAKTDHGAEIVYKSPVVSGDT SPRHLSNVSSTGSIDMVDSPQLATLADEVSASLAKQGL
- K27 MSSPGSPGTPGSRSRTPSLPTPPTREPKKVAVVRTPPKSPSSAKSRLQTAPVPMPDLKNVKSKIG STENLKHQPGGGKVQIVYKPVDLSKVTSKCGSLGNIHHKPGGGQVEVKSEKLDFKDRVQSKIGSL DNITHVPGGGNKKIETHKLTFRENAKAKTDHGAEIVY
- K16 MSSPGSPGTPGSRSRTPSLPTPPTREPKKVAVVRTPPKSPSSAKSRLQTAPVPMPDLKNVKSKIG STENLKHQPGGGKVQIINKKLDLSNVQSKCGSKDNIKHVPGGGSVQIVYKPVDLSKVTSKCGSLG NIHHKPGGGQVEVKSEKLDFKDRVQSKIGSLDNITHVPGGGNKKIE
- K25 MAEPRQEFEVMEDHAGTYGLGDRKDQGGYTMHQDQEGDTDAGLKAEEAGIGDTPSLEDEAAGHVT QARMVSKSKDGTGSDDKKAKGADGKTKIATPRGAAPPGQKGQANATRIPAKTPPAPKTPPSSGEP PKSGDRSGYSSPGSPGTPGSRSRTPSLPTPPTREPKKVAVVRTPPKSPSSAKSRL
- K32 MSSPGSPGTPGSRSRTPSLPTPPTREPKKVAVVRTPPKSPSSAKSRLQTAPVPMPDLKNVKSKIGS TENLKHQPGGGKVQIINKKLDLSNVQSKCGSKDNIKHVPGGGSVQIVYKPVDLSKVTSKCGSLGNI HHKPGGGQVEVKSEKLDFKDRVQSKIGSLDNITHVPGGGNKKIETHKLTFRENAKAKTDHGAEIVY
- K23 MAEPRQEFEVMEDHAGTYGLGDRKDQGGYTMHQDQEGDTDAGLKAEEAGIGDTPSLEDEAAGHVT QARMVSKSKDGTGSDDKKAKGADGKTKIATPRGAAPPGQKGQANATRIPAKTPPAPKTPPSSGEP PKSGDRSGYSSPGSPGTPGSRSRTPSLPTPPTREPKKVAVVRTPPKSPSSAKSRLTHKLTFRENA KAKTDHGAEIVYKSPVVSGDTSPRHLSNVSSTGSIDMVDSPQLATLADEVSASLAKQGL
- K44 MAEPRQEFEVMEDHAGTYGLGDRKDQGGYTMHQDQEGDTDAGLKAEEAGIGDTPSLEDEAAGHVT QARMVSKSKDGTGSDDKKAKGADGKTKIATPRGAAPPGQKGQANATRIPAKTPPAPKTPPSSGEP PKSGDRSGYSSPGSPGTPGSRSRTPSLPTPPTREPKKVAVVRTPPKSPSSAKSRLQTAPVPMPDL KNVKSKIGSTENLKHQPGGGKVQIVYKPVDLSKVTSKCGSLGNIHHKPGGGQVEVKSEKLDFKDR VQSKIGSLDNITHVPGGGNKKIE
- hTau23 MAEPRQEFEVMEDHAGTYGLGDRKDQGGYTMHQDQEGDTDAGLKAEEAGIGDTPSLEDEAAGHVT QARMVSKSKDGTGSDDKKAKGADGKTKIATPRGAAPPGQKGQANATRIPAKTPPAPKTPPSSGEP PKSGDRSGYSSPGSPGTPGSRSRTPSLPTPPTREPKKVAVVRTPPKSPSSAKSRLQTAPVPMPDL KNVKSKIGSTENLKHQPGGGKVQIVYKPVDLSKVTSKCGSLGNIHHKPGGGQVEVKSEKLDFKDR VQSKIGSLDNITHVPGGGNKKIETHKLTFRENAKAKTDHGAEIVYKSPVVSGDTSPRHLSNVSST GSIDMVDSPQLATLADEVSASLAKQGL
- hTau40 MAEPRQEFEVMEDHAGTYGLGDRKDQGGYTMHQDQEGDTDAGLKESPLQTPTEDGSEEPGSETSD AKSTPTAEDVTAPLVDEGAPGKQAAAQPHTEIPEGTTAEEAGIGDTPSLEDEAAGHVTQARMVSK SKDGTGSDDKKAKGADGKTKIATPRGAAPPGQKGQANATRIPAKTPPAPKTPPSSGEPPKSGDRS GYSSPGSPGTPGSRSRTPSLPTPPTREPKKVAVVRTPPKSPSSAKSRLQTAPVPMPDLKNVKSKI GSTENLKHQPGGGKVQIINKKLDLSNVQSKCGSKDNIKHVPGGGSVQIVYKPVDLSKVTSKCGSL GNIHHKPGGGQVEVKSEKLDFKDRVQSKIGSLDNITHVPGGGNKKIETHKLTFRENAKAKTDHGA EIVYKSPVVSGDTSPRHLSNVSSTGSIDMVDSPQLATLADEVSASLAKQGL

